# Automated analysis of large-scale NMR data generates metabolomic signatures and links them to candidate metabolites

**DOI:** 10.1101/613935

**Authors:** Bita Khalili, Mattia Tomasoni, Mirjam Mattei, Roger Mallol Parera, Reyhan Sonmez, Daniel Krefl, Rico Rueedi, Sven Bergmann

## Abstract

Identification of metabolites in large-scale ^1^H NMR data from human biofluids remains challenging due to the complexity of the spectra and their sensitivity to pH and ionic concentrations. In this work, we test the capacity of three analysis tools to extract metabolite signatures from 968 NMR profiles of human urine samples. Specifically, we studied sets of co-varying features derived from Principal Component Analysis (PCA), the Iterative Signature Algorithm (ISA) and Averaged Correlation Profiles (ACP), a new method we devised inspired by the STOCSY approach. We used our previously developed metabomatching method to match the sets generated by these algorithms to NMR spectra of individual metabolites available in public databases. Based on the number and quality of the matches we concluded that both ISA and ACP can robustly identify about a dozen metabolites, half of which were shared, while PCA did not produce any signatures with robust matches.

## Introduction

Nuclear Magnetic Resonance (NMR) spectroscopy is a powerful technique for metabolomic profiling. NMR does not consume the sample and has high accuracy and reproducibility. Single proton NMR spectroscopy (^1^H NMR) can be used to generate one-dimensional spectra of biofluids at high throughput and relatively low cost, facilitating the generation of large sets of spectral data.

A first step in NMR spectral analysis is usually to identify the main metabolites giving rise to a given spectrum or set of spectra. This is non-trivial, since human biofluids typically contain a large number of individual metabolites and their corresponding peak positions may overlap and are often affected by the pH, ionic strength and overall protein content of the fluid.

For small set of samples, expert analysis is therefore still the most accurate means for metabolite identification, yet for large collections of samples this approach is costly, time-consuming and potentially less reproducible. As a result, various methods have been suggested to assist or fully automate metabolite identification.

In their landmark paper on Statistical TOtal Correlation SpectroscopY (STOCSY), Cloarec et al. showed that analyzing the correlation patterns between features across a sizable collection of ^1^H NMR spectra has great potential for metabolite identification (Cloarec et al. 2005). This is because features corresponding to the same molecule (or molecules whose concentrations co-vary) tend to be significantly correlated in large datasets. Analyzing data from 612 mouse urine samples, they observed that the correlation matrix exhibited correlated peaks of features characteristic of valeramide, glucose, hippurate, 2-oxoglutarate, 3-hydroxyphenylpropionate, citrate, as well as methylamine, dimethylamine and trimethylamine.

Subsequent variations of STOCSY attempted to make clusters of NMR peaks to simplify the interpretation of large amount of information stored in the correlation matrix from STOCSY analysis. Recoupled-STOCSY (R-STOCSY) employs a variable size bucketing method to reduce the dimensionality of NMR data and statistical recoupling of variables (SRV) to identify correlations between distant clusters (Blaise et al. 2010). Iterative-STOCSY (I-STOCSY) aims at separating the inter-metabolite connections from intra-metabolite connection by recursively applying STOCSY analysis first from a selected “driver peak” and then for all peaks correlating with the driver peak above some threshold (Sands et al. 2011). SubseT Optimization by Reference Matching (STORM) selects subsets of ^1^H NMR spectra that contain specific spectroscopic signatures of biomarkers differentiating between different human populations (Posma et al. 2012).

Once metabolite identification has been achieved, the next challenge is to quantify metabolite concentrations. This process works robustly only for a relatively small set of metabolites, and requires expert refinement when using publicly available quantification tools such as BATMAN (Hao et al. 2012), FOCUS (Alonso et al. 2014), BAYESIL (Ravanbakhsh et al. 2015), ASICS (Tardivel et al. 2017), AQuA (Röhnisch et al. 2018) or rDolphin (Cañueto et al. 2018). This is unsatisfactory in light of the fact that a sizable number of metabolites have been identified in human biofluids. For example, the latest version of the Human Metabolome Database (HMDB 4.0) (Wishart et al. 2018) includes more than 1’500 metabolites with ^1^H NMR spectra, 179 of which have been identified in urine (Bouatra et al. 2013) and 67 in serum (Psychogios et al. 2011; Nagana Gowda et al. 2015).

There are several reasons why it remains difficult to perform fully automated quantification from large-scale NMR data for the vast majority of metabolites: First, human biofluids typically contain a large number of individual metabolites whose concentrations vary across several orders of magnitude. This makes it difficult to disentangle the contributions of metabolites with low concentrations, in particular when their NMR features are not unique. Second, the exact feature positions depend on the biofluid and may have been different when acquiring reference spectra. Third, while the number of reference spectra continues to grow, they are certainly not yet exhaustive.

In two recent studies we showed that the limitations of targeted NMR metabolomics can be overcome to some extent if the goal is to link metabolites to external variables (Rueedi et al. 2014; Rueedi et al. 2017). Specifically, in the context of genome-wide association studies (GWAS) applied to metabolomics (known as mGWAS) the aim is to associate metabolites with genotypic variants. We observed that the effect of a genetic variant on the concentration of a metabolite often translates into associations with all or many features of the metabolite NMR spectrum. The set of association scores with all measured features provides a *pseudospectrum* across the full range of ppm covered by ^1^H NMR spectra. The challenge is then to identify the metabolite underlying the most significant associations. To this end we developed the analysis tool *metabomatching*, which takes as input a pseudospectrum and a collection of reference spectra for individual metabolites such as found in HMDB (Wishart et al. 2018). Our previous work showed that metabomatching works well to prioritize the most likely metabolite candidates for pseudospectra derived from metabolic feature association with genotypes (Rueedi et al. 2014; Rueedi et al. 2017; Raffler et al. 2015).

In the present work we show that the metabomatching methodology can also be used for identifying metabolites that vary across large collection of samples without the need for any external variables associated with this variation. We investigate three methods that identify co-varying spectral features within large-scale NMR data. Specifically, we compare Principal Component Analysis (PCA), the Iterative Signature Algorithm (ISA) and averaged correlation profiles (ACP) inspired by the STOCSY approach. For each method, we devise a principled way for processing their output into pseudospectra. We then evaluate systematically in which cases metabomatching provides strong evidence for matching metabolites, and assess to what extent the three methods provide consistent or complementary output. Our analysis software for the unsupervised generation of metabolomic signatures from large-scale NMR data and integration with metabomatching (including further documentation) is available at https://github.com/BergmannLab/metabomodules.

## Analysis & Results

### Metabomatching

Our original metabomatching method was designed to match the NMR spectra of individual metabolites recorded in a database with association profiles between metabolome features and an external variable, typically a SNP genotype. Specifically, for a metabolite *m*, metabomatching computes the sum

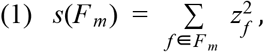

where *F*_*m*_ is the set of *N*_*f*_ features that fall within a neighbourhood of a peak of *m* according to the database, and *z*_*f*_ denotes the significance value for feature *f*, and is given by 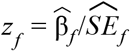, where 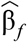 and 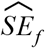 refer to the point estimates of the effect size and its standard error, respectively. Under the null hypothesis of normally and independently distributed *z*_*f*_, the sum *s* follows a χ^2^ distribution with *N*_*f*_ degrees of freedom, and metabomatching defines the score for *m* as the negative logarithm of the nominal p-value for the sum. This score is then used to rank all tested metabolites: the higher the score, the better the match to the metabolite.

In this work, we show that metabomatching also works with pseudospectra capturing the internal structure of large-scale NMR data rather than their correlation with external variables. Specifically, our premise is that in sizable sample collections there is sufficient power for methods identifying coherent features that may point to the same metabolite. Since such methods typically provide their output in terms of correlations or *z*-scores, or can be transformed into either of them, we extended metabomatching to work also with such inputs (see Methods for details).

We used three methods for identifying weighted sets of co-varying spectral features from large-scale NMR data that can be used as input for metabomatching (see Methods for more details and Figure 1 for illustration of the workflow).

**Figure 1:**
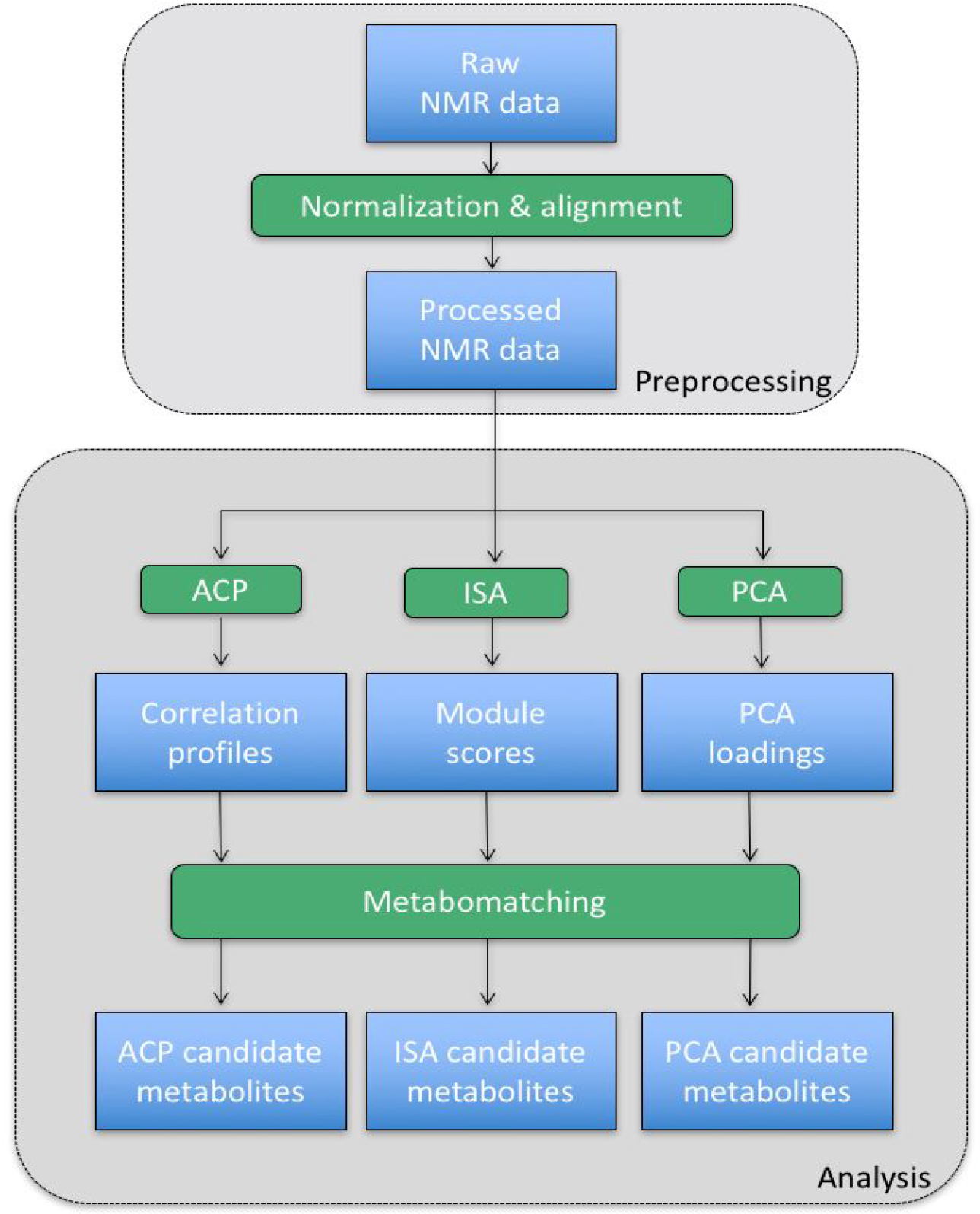
Workflow for unsupervised analysis of large-scale NMR data. Raw ^1^H NMR data are normalized then aligned. These processed profiles are used as input for the Averaged Correlation Profile (ACP), Iterative Signature Algorithm (ISA) and Principal Component Analysis (PCA) methods, which output correlation profiles, module scores and PCA loadings, respectively. These outputs constitute possible pseudospectra for metabomatching which identifies the most plausible candidate metabolites underlying the coherent feature variations.

### Correlation-based pseudospectra

Our first approach to select co-varying features was to use the correlations between features across all samples. We faced two challenges: First, using the correlations of a given feature with all features as a pseudospectrum would break the scoring algorithm in metabomatching, because the correlation of a feature with itself is one, corresponding to an infinite *z*-score. Second, we wanted a relatively small set of pseudospectra as input for metabomatching, so we needed to rank possible input sets to select only the most relevant.

To address these challenges we devised an algorithm that ranks all pairs of sufficiently distant features and computes averaged correlation profiles (ACP) for pairs of highly correlated features. Strictly speaking ACPs do not consist of correlations, but for highly correlated feature pairs the correlations to other features tend to be very similar, while none of the averages equals to one. Within metabomatching these ACPs are then translated into z-scores using the Fisher z-transformation (see Methods for more details). We also tried hierarchical clustering to define pseudospectra from multiple highly similar features, but this approach did not work as well (see Methods).

### Iterative Signature Algorithm

The ISA has been designed for the unsupervised identification of coherent subsets in large-scale data (Ihmels et al. 2004; Bergmann et al. 2003). Specifically, coherence between features is *not* defined by total correlation across all samples, but rather for a subset of samples for which a set of features takes more extreme values than for the rest of the samples. A heuristic iterative procedure starting with random features is used to refine so-called *modules*, consisting of such self-consistent subsets of features *and* samples. Each module is defined for a set of two thresholds, determining how extreme the features and samples are allowed to be. Importantly, scanning through an array of thresholds usually identifies a relatively small set of modules (or very similar families thereof), i.e. much smaller than the number of samples or features.

In order to obtain a pseudospectrum for each module we averaged each feature (whether part of the module or not) across the samples assigned to the module. From these averages we computed z-scores. By definition the features of the module have the most extreme z-scores, yet other features that were just below the threshold may also have a sizable contribution. We used these z-scores as input for metabomatching. Specifically, since we allowed feature scores with positive and negative signs we used the plus/minus mode of metabomatching. This allows for detecting metabolites corresponding either to the negative or positive features (see Methods for more details).

### Principal Component Analysis

We also used PCA to compute the loadings of all features onto the eigenvectors of the sample-sample correlation matrix across all features. These eigenvectors (or eigen-samples) characterise independent axes of variation in sample space. The corresponding eigenvalues reflect the fraction of total variation explained by each eigenvector. It was not clear a priori which principal components might characterize variation due to single metabolites. We therefore applied metabomatching to all of them. Specifically, we generated pseudospectra by standardizing the loadings corresponding to each eigen-sample (see Methods for more details).

### Many pseudospectra defined by the ACP method and ISA match to urine metabolites

We observed a trend of elevated metabomatching scores for pseudospectra corresponding to principal components with small eigenvalues (starting from component #505), jointly explaining only 1% of variation (Supplementary Figure 1A). The last nine principal components matched to hippurate, but disappeared when running PCA on the metabolome stripped of highly correlated features (see Methods; Supplementary Figure 1D and 1F). Additionally, the adjusted scores of all potential hits decreased significantly for the stripped metabolome (Supplementary Figure 1E). We therefore concluded that PCA is not well-suited for generating robust metabolite signatures.

In contrast, our ACP method and ISA resulted in a sizable number of pseudospectra for which metabomatching produced robust matches to urine metabolites (see Figure 2 and Methods for details). Specifically, both ACP and ISA identified feature sets pointing to glucose, citrate, ethanol, hippurate and P-hydroxyphenylacetate (Supplementary Figures 2-11). Glucose and hippurate were amongst the metabolites associated by Cloarec et al. with the correlation matrix of NMR data obtained from mouse urine samples (Cloarec et al. 2005).

**Figure 2:**
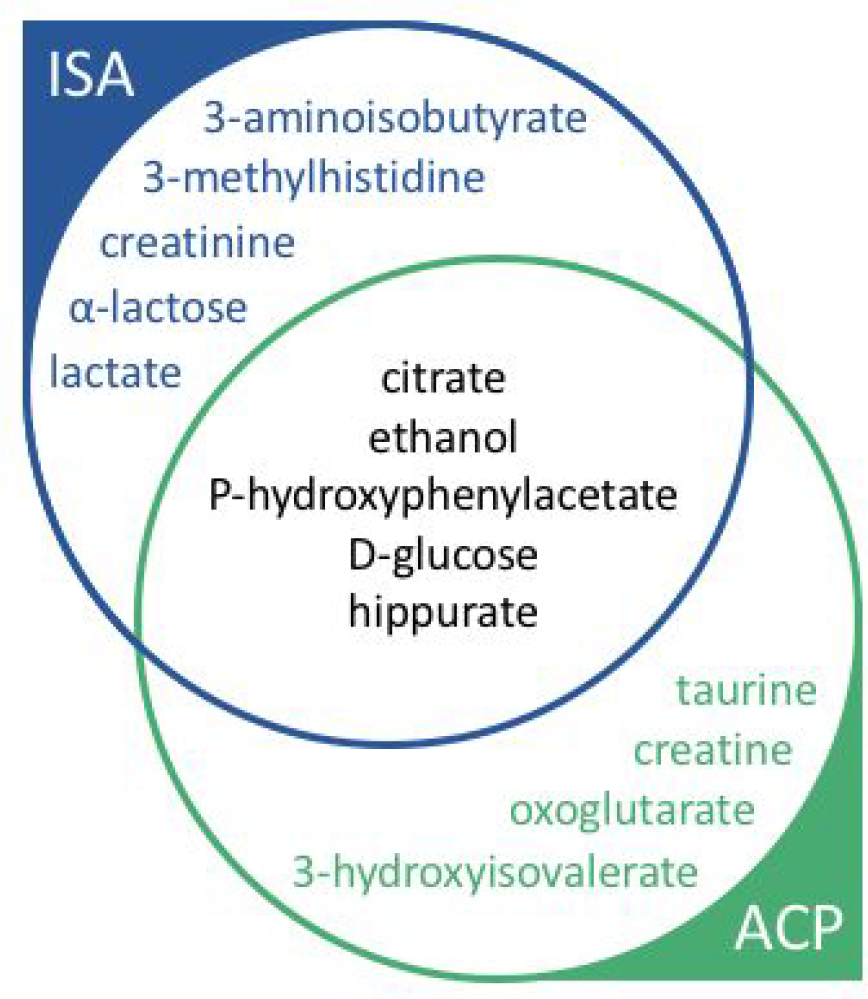
Urine metabolites that were robustly matched by metabomatching to pseudospectra derived from average correlation profiles (ACP, green), the Iterative Signature Algorithm (ISA, blue) or both methods (black).

P-hydroxyphenylacetate shares an aromatic ring with 3-hydroxyphenylpropionate, another metabolite highlighted in the original STOCSY paper (Cloarec et al. 2005). Both compounds are part of the phenylalaline metabolism and occur as products of bacterial degradation of aromatic compounds. In human urine high concentrations of these compounds may reflect an overgrown Clostridium species in gut microbiota which has been associated with autism spectrum disorders (Xiong et al. 2016).

In healthy humans urine glucose should be very low, but concentrations may be elevated due to diabetes or chronic kidney disease (CKD), conditions which are prevalent in the CoLaus population from which we obtained the urine samples.

Citrate is an additive commonly used by the food industry and it is also synthesised as an intermediate product in the tricarboxylic acid cycle, a central pathway that releases stored energy from fat, proteins and carbohydrates. Low urinary citrate is associated with CKD and kidney stone formation.

There are a number of metabolites that could only be picked up robustly with one of the two methods: With ISA we found modules matching 3-aminoisobutyrate (an end product of nucleic acid metabolism that has been considered a potential biochemical marker for cancer (Nielsen & Killmann 1983); Supplementary Figure 12), creatinine (a breakdown product of creatine, whose high and relatively stable concentration in urine is often used for normalisation; Supplementary Figure 13), as well as the milk sugar lactose (Supplementary Figure 14) and its bacterial breakdown product lactic acid (Supplementary Figure 15). Conversely, the ACP method picked up taurine (2-aminoethanesulfonic acid; Supplementary Figure 16), an organic compound widely distributed in animal tissues and a major constituent of bile; creatine (Supplementary Figure 17); oxoglutarate (α-ketoglutarate; Supplementary Figure 18), an important biological compound produced by deamination of glutamate, and an intermediate in the Krebs cycle; and 3-hydroxyisovalerate (Supplementary Figure 19), which is a byproduct of valine, leucine and isoleucine degradation and a marker for biotin deficiency (Ziegler et al. 1996). All these compounds have in common that they can occur at relatively high concentrations in human urine.

Correcting features for significant covariates, both methods also picked up carnitine, which owes its name to its high concentration in meat (see Methods for more details). While it is produced in both animal and plant cells, this may explain why we could detects its signature in the human data.

### Metabolite concentration pseudo-quantification with NMR features of matched pseudospectra

We next investigated whether the sets of weighted NMR features generated by ISA or the ACP method can not only be used to identify metabolites, but also facilitate their quantification. Specifically, we hypothesized that the leading features suggested by our algorithms for a certain metabolite may be better suited for quantification than using all features of reference spectra from public databases like UMDB. There are two possible reasons for this. First, the exact feature positions extracted from the data by ISA or the ACP method may be more accurate, even if most of them match those of the reference spectrum within the margin of the matching window. Second, for metabolites with several peaks, only a subset might have been picked up by these algorithms. Indeed, both ISA and the ACP method may leave out peaks which did not contribute coherently to the peak set since their signal was too noisy (e.g. due to overlap with those from other metabolites).

To test this, we used discrete integration to estimate metabolite concentrations, according to

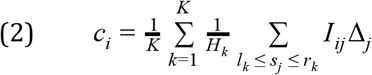

where *K* is the number of multipets, *H*_*k*_ the number of protons in multiplet *k*, [*l*_*k*_, *r*_*k*_] the range of multiplet *k*, *s*_*j*_ the chemical shift of feature *j*, *I*_*ij*_ the intensity of this feature in individual *i*, and *Δ*_*j*_ the width of the bin at *s*_*j*_. For example, for hippurate *K* = 4, *H* = [2,2,1,2], *l* = *m*-0.025, *r* = *m*+0.025, where *m* = [3.98, 7.54, 7.65, 7.84]. We then evaluated this concentration first using the peak positions from the reference spectrum as listed in UMDB, and second using the peak positions as suggested by the pseudospectrum found by the modular approach.

We applied the above pseudo-quantification methods to glucose and ethanol, for which we had other relevant measures. For urine glucose the most relevant independently measured biomarkers was fasting blood glucose. For urine ethanol, relevant biomarkers included serum asialotransferrin and disialotransferrin, which combined are known as carbohydrate-deficient transferrin (CDT) which is a biomarker for heavy alcohol use. Furthermore, self-reported alcohol consumption was available. These biomarkers were measured in a different biofluid (i.e. blood), which was collected on the same day as the urine sample.. We argue that detecting significant correlations between our estimated metabolite concentrations and these biomarkers provides a proof of concept that our quantification is reliable, and comparing correlations between different quantification approaches provides a means to evaluate them.

The ^1^H NMR spectra of glucose has nine multiplets. Including all these multipets chemical shifts from the UMDB database (Table 1) to perform pseudo-quantification based on equation (2), we obtain a correlation of 0.46 (with a 95% confidence interval (CI) of [0.41, 0.52]) between the pseudo-quantification and fasting blood glucose. This correlation increases to 0.50 [0.44, 0.55] if the subset of seven multiplets from ISA module #16 (Figure 3A) is used for pseudo-quantification (see Methods for details). From 179 ACP pseudospectra, two robustly matched glucose, one from averaging the correlation profiles from feature pair 3.48 & 5.24 and another from averaging correlation profiles from 3.89 & 5.24 (Figure 3B and 3C), each capturing four multiplets out of nine multiplets of glucose (Table 1). Using the subsets of the four peaks from 3.48 & 5.24 we obtain a correlation of 0.48 [0.43, 0.54] between glucose pseudo-quantification and fasting blood glucose while using the subsets of four peaks from 3.89 & 5.24 a correlation of 0.44 [0.38, 0.49] was obtained.

**Table 1:** Correlation between pseudo-quantification and measured biomarkers of glucose and ethanol.

| **Urine Metabolite** | **Feature source** | **Multiplet positions (ppm)** | **Related bio-marker** | **Correlation [95% CI]** |
| --- | --- | --- | --- | --- |
| glucose | UMDB | 3.23, 3.40, 3.46, 3.52, 3.73, 3.82, 3.88, 4.63, 5.22 | serum glucose | 0.46 [0.41, 0.52] |
| glucose | ACP: 3.48 & 5.24 | 3.40, 3.48, 4.65, 5.24 | serum glucose | 0.48 [0.43, 0.54] |
| glucose | ACP: 3.89 & 5.24 | 3.82, 3.89, 4.65, 5.24 | serum glucose | 0.44 [0.38, 0.49] |
| glucose | ISA: module #16 | 3.25, 3.41, 3.48, 3.50, 3.89, 4.65, 5.24 | serum glucose | 0.50 [0.44, 0.55] |
| ethanol | UMDB | 1.17, 3.65 | serum CDT | 0.29 [0.23, 0.35] |
| ethanol | ACP: 1.18 & 3.67<br>ISA: module #57 | 1.18, 3.67 | serum CDT | 0.16 [0.10, 0.22] |
| EtG | (Nicholas et al. 2006) | 1.24, 3.30, 3.52, 3.71, 3.99, 4.48 | serum CDT | 0.36 [0.30, 0.42] |
| EtG | ISA: module #240 (matching subset) | 1.24, 3.52, 4.47 | serum CDT | 0.46 [0.40, 0.51] |

**Figure 3:**
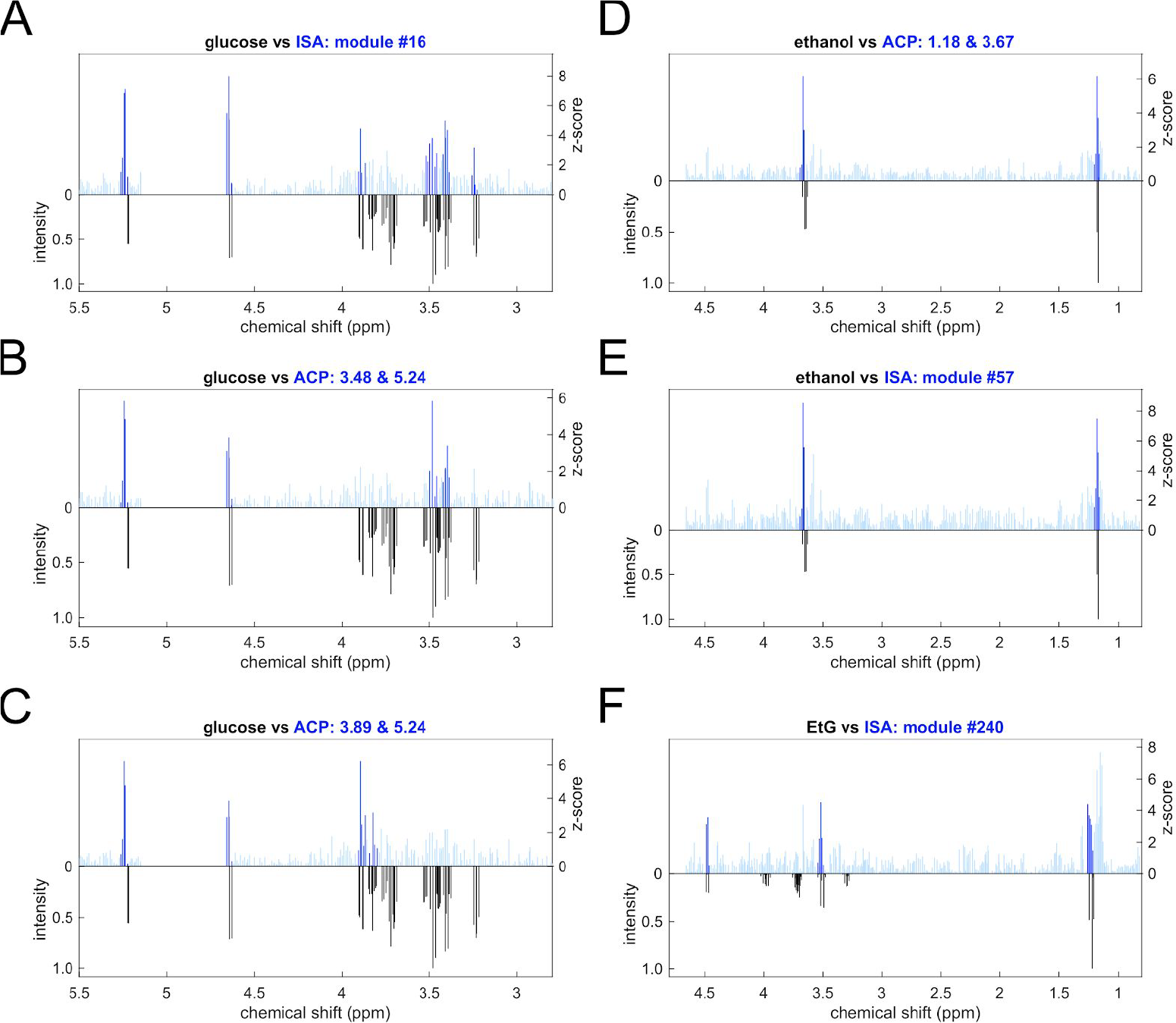
Pseudospectra from ACP and ISA algorithms matching glucose, ethanol and EtG. Each plot shows the pseudospectrum in blue in the upper half and the reference spectrum from UMDB in black and in the lower half. Dark blue indicates chemical shifts and their +/−0.025 ppm vicinity that were used for pseudo-quantification.

Combining the peak subsets from both pseudospectra to a set of 6 peaks did not improve the correlation beyond 0.48 [0.43, 0.54] (see Table 1 and Methods for more details on peak sets used for pseudo-quantification).

The ^1^H NMR spectra of ethanol only has two multiplets (as well as one singlet from the hydroxyl group, which cannot be discerned in water solutions, like urine). Using the ppm positions of these two multiplets from the UMDB database to carry out pseudo-quantification, we observed a relatively low correlation of 0.29 [0.23, 0.35] between the quantification and CDT levels in serum (Table 1, see Supplementary Table 1 for other alcohol markers). The ACP method identifies one module, 1.18 & 3.67, and ISA identifies two modules (#57 and #240) that metabomatching matched to ethanol (Supplementary Figures 6,7 and 22). The positions of the ACP peak set (i.e. 1.18 & 3.67) were identical to those of ISA module #57 and were more similar to the ethanol spectrum (achieving a higher adjusted score in metabomatching) than those of ISA module #240 (Figure 3D-3F). Nevertheless, pseudo-quantification with equation (2) using these positions yielded a lower correlation of 0.16 [0.10, 0.22] with CDT levels than the UMDB reference peaks (0.29 [0.23, 0.35]) (Table 1). This is due to a high correlation of CDT with the features at 1.145-1.155 ppm which are within a 0.025 ppm neighborhood of the UMDB ethanol peak at 1.17 ppm but not within the same neighborhood of the 1.18 ethanol peak from ACP and ISA module #57 (Supplementary Figure 20). Yet, these peaks at 1.145-1.155 ppm are unlikely to correspond to ethanol, since their correlation with the other ethanol peak at 3.67 ppm is much weaker than the correlation between the 1.18 ppm and 3.67 ppm peaks, and they may be linked to a different metabolite whose concentration is correlated with CDT (Supplementary Figure 21). In contrast, summing up the intensities over all the features of ISA module #240 with a *z*-score above 3 as a quantification measure (in the absence of any multiplet information), we found that this measure had a correlation of 0.51 [0.46, 0.57] with the CDT measurements.

To better understand why module #240 correlates more strongly to the alcohol consumption biomarker while being a worse match to ethanol than module #57 (Figure 3), we studied whether any of its features point to other compounds related to ethanol metabolism. Indeed, we found that this module contains three features, at 1.26 ppm, 3.52 ppm and 4.47 ppm, that individually achieve higher correlations to CDT (0.40 [0.34, 0.45], 0.29 [0.23, 0.35], 0.33 [0.27, 0.39] respectively) than the features mapping to ethanol. Interestingly, these features appeared to be close to those of ethyl glucuronide (EtG), a direct product of ethanol non-oxidative metabolism by conjugation with uridine diphosphate (UDP)-glucuronic acid, which had previously been detected in ^1^H NMR spectra of liver extracts (Nicholas et al. 2006) and more recently in human urine of alcohol drinkers (Kim et al. 2013). To confirm EtG as a possible match for ISA module #240, we added its features as extracted from (Nicholas et al. 2006) to the metabomatching library manually, since EtG had no entry in UMDB. We observed that ethanol and EtG spectra together provided a better match to ISA module #240 than ethanol alone (Supplementary Figure 22). Interestingly, the distance between the two peaks corresponding to the doublet at 4.48 ppm is about 0.0126 ppm corresponding to a coupling of 8.8 Hz (for a 700 MHz spectrometer) consistent with the coupling of 8 Hz reported in (Nicholas et al. 2006) (see Supplementary Figure 23 and Supplementary Information for more details).

Performing the pseudo-quantification of EtG using either a peak set of 6 reference positions extracted from (Nicholas et al. 2006), or a subset of 3 features from module #240 resulted in a correlation with CDT levels of 0.36 [0.30, 0.42] for the former and 0.46 [0.40, 0.51], for the latter (Table 1). This indicates that EtG correlates better with CDT than ethanol, which is in agreement with the fact that EtG is detectable for a longer time window (2-5 days) than ethanol (12-24 hours) in urine, and CDT is a marker for heavy alcohol use (at least five drinks a day over a period of two weeks before giving the sample (Solomons 2012)). Remarkably, the pseudo-quantification facilitated by the three features of module #240 correlates even more strongly with CDT than the full set of EtG reference features, presumably because these features have the best signal to noise ratio and optimal position for our data. They may therefore constitute a promising urine biomarker for heavy alcohol consumption. Indeed, while the correlation between EtG pseudo-quantification and CDT measure increases to 0.59 [0.46, 0.72] focusing on subjects who have self-reported heavy drinking, pseudo-quantification of module #240 gives rise to a slightly higher correlation of 0.61 [0.48, 0.74].

## Conclusions and discussion

In this work, we developed and tested new methodologies for analysing large-scale ^1^H NMR spectroscopy data. Building on previous ideas to use the correlation structure of such data to generate metabolic signatures, we investigated three complementary methods for generating such signatures and benchmarked the methods in terms of how many of their signatures could be matched with reference spectra in public databases. By design, these approaches will only identify metabolites with at least two distinct peaks, and therefore complement peak-picking identification approaches which tend to focus on single peak metabolites.

We found that average correlation profiles (ACP) of highly correlated feature pairs, a method inspired by STOCSY, as well as the Iterative Signature Algorithm (ISA), which was first developed for modular analysis of gene expression data, both identified about a dozen of metabolites robustly, half of which were shared. In contrast, Principal Component Analysis (PCA) did not generate any pseudospectra with robust metabomatching, likely because leading components explain variation driven by many metabolites.

While ACP is designed to pick up individual metabolites with at least two (non-proximal) features in their spectrum (or those of metabolite pairs whose concentrations are coupled), ISA is able to generate modules where many features exhibit coherent variation, yet potentially only over a subset of samples. We believe that this may be particularly useful when integrating data from a heterogeneous set of samples (e.g. including those from diseased or medicated subpopulations).

An interesting property of our modular approach is the fact that the feature sets identified by ACP or ISA do not need to match perfectly with those of the reference spectrum of the corresponding compound. Indeed, the two ACP signatures matching glucose each only cover four and jointly six of the nine glucose peaks, while the ISA module with the best match to glucose includes seven of its peaks. Adding the “missing” peaks in our quantification slightly reduced the correlation with serum glucose, indicating there is a marginal improvement in the pseudo-quantification using ppm positions only from the multiplets found by our algorithms rather than the database. Further work will be needed to substantiate this observation.

Another interesting aspect of our approach is that modular feature sets may match multiple compounds. Our current implementation of metabomatching allows simultaneous identification of up to two compounds. Indeed, our finding that ISA picked up a module whose signature mapped well to ethanol and its specific metabolic product ethyl glucuronide demonstrated the potential power of the ISA to identify metabolite pairs within the same pathway. Moreover, the strong observed correlation of this module with the alcohol abuse marker CDT was likely driven by the fact that ISA can extract context specific covariance, which in this case is strongest in samples with particularly high alcohol consumption. This module also highlighted that using the relevant chemical shifts found by the module rather than all shifts from the reference database can lead to more accurate quantification of the underlying metabolites, due to different contribution of shifts specific to the experimental conditions in the complex urine spectra.

Extending metabomatching beyond compound pairs is challenging due to the large number of possible triplets, let alone higher order combinations, but could be feasible in future work when taking a targeted approach, which uses pathway knowledge to scan interesting metabolite combinations.

A critical ingredient to our analysis was to transform the signatures generated by the different methods into a universal format (i.e. z-scores) as input for our metabomatching tool that we had developed previously for the analysis of feature signatures generated by regression on external variables. Indeed, being able to query both internal and external signatures of large-scale NMR data against a reference dataset of known spectra from individual metabolites is pivotal for exploring new methods dissecting the auto-as well as cross-correlation structure for integrative analyses.

In conclusion, we believe that our study using fewer than 1000 samples gives ample evidence for the potential of modular analysis of large-scale NMR data, and that increased sample sizes are likely to results in further identifications and more accurate quantifications of individual metabolites. Importantly, to this end all our analysis software is made publicly available on github (https://github.com/BergmannLab/metabomodules).

## Methods

### Preprocessing

For this study, we used 968 ^1^H NMR spectra acquired in urine samples from the CoLaus cohort (Rueedi et al. 2014). The samples are referenced according to the TSP signal, phase-corrected, baseline-corrected and binned into 23,488 bins.

We aligned and rebinned the spectra using the FOCUS tool (Alonso et al. 2014) choosing its parameters such that the resulting data still had a relatively high resolution (<0.02 ppm) and a large number of 687 peaks. We set the downsampling frequency parameter *window.fs* to 1 (indicating no downsampling), the sliding window length *window.length* for spectral segmentation to 0.03 ppm, the minimum peak width parameter *peak.DFL* to 0.02 ppm and the peak sample frequency parameter *peak.pS* to 0.2, while keeping the rest of parameters at their default values.

We then normalized the FOCUS data: applying a log10-transformation, then standardizing across features (effectively normalizing the concentration of each sample), then standardizing across samples (making feature intensities comparable).

### Confounding

The main confounding factors of the NMR data that we investigated here are age, sex, serum creatinine and urine creatinine ((Rueedi et al. 2014; Rueedi et al. 2017)). In order to allow for the identification of metabolites that may be hidden by confounding, we generated a dataset of residuals, created by regressing out the confounders from the feature metabolome.

### Metabomatching

In addition to pseudospectra from regression analysis, still provided as columns headed by *beta*, *se*, and *p*, we extended metabomatching to accept pseudospectra produced by PCA, ACP, and ISA as columns headed by *pca*, *cr*, and *isa* respectively. For ACP pseudospectra, we translate a correlation *c* to a *z*-score with the Fisher transformation *z* = λ *arctanh*(*c*). For independent features, 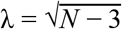 gives *z*-scores with unit standard deviation, when *N* is the number of samples across which the correlations are computed. However, since proximal features are usually not independent we allow for a user-provided estimate for λ (obtained from the pairwise feature-feature correlations matrix), or re-estimate λ from the given correlations.

For ISA and PCA pseudospectra, we standardize the loadings or module scores. We also introduced a measure of the quality of a match, which allows, in particular, the comparison of matches between different pseudospectra. This *adjusted score* is obtained by reshuffling the pseudospectrum, and defining a heuristic *p*-value by the number *N*_*p*_ of all *N*_*r*_ reshuffled pseudospectra that produces a metabomatching score (for any reference spectrum) higher than the the metabomatching score for the input pseudospectrum of the highest ranked reference spectrum. This *p*-value is defined as (*N*_*p*_ + 1)/(*N*_*r*_ + 1), and the adjusted score as − *log*(*p*). We used *N*_*r*_ = 999 which sets the highest possible adjusted score to 4.

We took special care in our feature reshuffling procedure to obtain a conservative adjusted score. Specifically, metabomatching identifies *cut points* in the pseudospectrum that separate it into peak-preserving clusters of features and only re-shuffles these clusters. The cut points are obtained as follows. Let *f*_*i*_ be the positions on the chemical shift axis of the metabolome features, sorted such that *f*_*i*_ < *f*_*i* +1_, and *C* the set of cut points. Metabomatching first populates *C* with all features which were separated by more than δ_*gap*_ from their predecessor (i.e. *f*_*i*_ > *f*_*i*−1_ + δ_*gap*_). δ_*gap*_ = 0.3 ppm is used as default value. Next it sorts the remaining features by their corresponding absolute-valued z-scores. Starting from the feature with the lowest absolute z-score it adds features to *C* provided they have a distance greater than δ_*min*_ to any features already assigned to *C* and an absolute z-score below a threshold *z*_*min*_. Default values are 0.04 *ppm* for δ_*min*_ and the standard deviation of all absolute z-scores across the features of a given pseudospectrum for *z*_*min*_.

### ACP: Averaged Correlation Profile

We devised the following greedy approach to populate a list *L* of feature pairs and their corresponding correlation profiles *c* as input for metabomatching: (1) We compute and sort all pairwise correlations *C*_*ij*_ between features *f*_*i*_ and *f*_*j*_ separated by at least 0.1 *ppm* across all samples. (2) Starting with the feature pair *P* = (*i*, *j*) with the highest correlation we iteratively add it to *L* unless there is already a feature pair in *L* whose features are in close proximity (i.e. within 0.1 *ppm*) to *f*_*i*_ and *f*_*j*_, respectively. The pseudo-correlation profile is defined as the average of the correlation profiles of *f*_*i*_ and *f*_*j*_: *c*_*k*_ = (*C*_*ik*_ + *C*_*jk*_)/2. The correlation profiles of strongly correlated features typically are very similar to each other. Consequently their average is close to both of them, while avoiding any elements equal to one (as *c*_*i*_ = *c*_*j*_ = (1 + *C*_*ij*_)/2 < 1 provided that *C*_*ij*_ < 1).

As an alternative approach we tried agglomerative clustering of features iteratively joining features (or sets of features) whose correlation was above a threshold Cmin. Each time we joined (sets of) features we computed a new meta-feature as their average and its correlation to all remaining (meta-)features. For each cluster of features we then computed the mean correlation profile.

### ISA: Iterative Signature Algorithm

We first ran the ISA algorithm to generate modules from the NMR data using the default values for the parameters except the following: both row and column thresholds were changed from the default values {1,2,3} to {1,2,3,4,5,6} to produce modules containing fewer rows or columns that are more likely to represent single metabolites, the number of seeds was increased from 100 to 250, and the correlation threshold below which ISA considers two modules equal was lowered from 0.95 to 0.50 to favour diversity in modules.

We allowed feature scores to be either positive or negative, while sample scores were always positive. This means that modules can include features which have on average higher or lower intensities in the selected samples than for the remaining samples. Modules which include a mixture of positive and negative are likely to represent (at least) two metabolites whose concentrations are inversely related to each other (like for a substrate and its product).

This procedure generated 216 modules. To select the 179 modules to pass to metabomatching, we sorted them by the size of their basin of attraction. To measure the basin size, we ran ISA a second time with the same parameters, but on 10,000 seeds, turning off sweeping, and keeping all converged modules (by setting the purge option to false). We then assigned a run 2 module to the basin of the run 1 module with which it has the highest correlation (provided that correlation be greater than 0.5). We assumed the run 1 modules attractor basin size is approximately equal to the count of modules from run 2 which were assigned to them in the previous step. Finally we passed to metabomatching the 179 modules from run 1 with largest basins.

### PCA: Principal Component Analysis

We used the sklearn library for Python (2.7) to compute all principal components of the preprocessed data. Specifically, we used the *decomposition.PCA* object, with *n_components* set to 687 and *svd_solver* set to full.

### Identification of metabolites

After running metabomatching on all the pseudospectra generated from our modular algorithms ISA, ACP and PCA, we applied a filtering algorithm to select only the most robust matches among all pseudospectra. The filtering passes only those pseudospectra which achieve an adjusted score above 2 with their top metabolite match (ensuring the pseudospectra finds a reasonable match by metabomatching) and have at least one peak with z-score above 4 (ensuring there is a strong signal).

Out of 179 pseudospectra from ACP, 10 metabolite signatures pass the filtering. Amongst these, only one pseudospectrum, i.e. 3.87 & 3.75, the top match to hydroxypropionate (Supplementary Figure 24) did not look convincing because of the only slightly lower scored match to mannitol and a reasonable match to arabitol (Supplementary Figure 24).

Among 179 pseudospectra from the ISA modules, 19 metabolite signatures passed the filtering. Amongst these we discarded 9 for one or more of the following three reasons: (1) There are several metabolites which all achieve high adjusted scores (Supplementary Figures 25 and 26). (2) There is at least one strong peak in the pseudospectrum that does not match with any of the spectra of the best matching metabolites (see Supplementary Figures 27-32). This may happen for pseudospectra with a large number of peaks. These pseudospectra are not necessarily biologically irrelevant and might carry a signature for two or more related metabolites (like the aforementioned module #240 with features from both ethanol and EtG). (3) Some pseudospectra passing the filter, nevertheless did not appear to have sufficiently strong signals at the peak positions of their putative matching metabolites (Supplementary Figures 33 and 34).

We analysed all 687 pseudospectra from principal components with metabomatching and only 5 metabolite signatures passed the default filtering (Supplementary Figures 35-39). We observed an elevation in metabomatching scores for the last principal components (all above component #505 which jointly explain only 1% of variation; Supplementary Figure 1A). The last 9 principal components all matched hippurate (Supplementary Figure 1D). Yet, after removing one feature from all feature pairs that correlated above 0.95 (i.e. eight features from 34 feature pairs all belonging to 7.54-7.56, 7.63-7.65, 7.83-7.84 and 3.96 ppm regions matching the four hippurate multiplets). Running metabomatching on all pseudospectra generated from the principal components of the remaining features, only five passed the filtering (Supplementary Figure 1F). None of these seemed convincing when applying the same criteria as for the ISA modules.

### Pseudo-quantification of metabolite concentrations

To perform the pseudo-quantification based on the pseudospectra of the modules, we defined a set of multiplet positions to use in equation (2) for each module that robustly identified a metabolite. This set is composed of all the chemical shifts from the module of interest with z-scores above 3 and within 0.025 ppm of the multiplet positions of the matching metabolite in reference database. It includes all relevant peaks from the metabolite detected by the modular analysis. Table 1 contains all the multiplets for glucose, ethanol and EtG specified in reference database and sets of chemical shifts selected by ACP and ISA.

## Author contributions

SB and RR designed the study. The manuscript was written by BK and SB. The correlation-based methods was implemented and applied by BK. The modular analysis with ISA was performed by MM and RR. PCA was run by MT, DK and BK. All authors discussed the results and implications, and contributed to the manuscript.

## Acknowledgments

This work was supported by the Swiss National Science Foundation (grant FN 310030_152724/1) and the NIH (grant R03 CA211815).

## Software availability

https://github.com/BergmannLab/metabomodules

