## Supplementary Materials for "Automated analysis of large-scale NMR data generates metabolomic signatures and links them to candidate metabolites"

[Y]Co-last author

### Supplementary Information

#### *Analyzing coupling constant of EtG doublet at 4.48 ppm*

We measured the gap between the two peaks from ISA module #240 around 4.48 ppm with the highest z-scores. These peaks correspond to two consecutive bins at 4.4691ppm and 4.4817 ppm in the aligned dataset using FOCUS (Alonso et al. 2014) (c.f. preprocessing section in Methods). This gap is equal to 0.0126 ppm equivalent to 8.82 Hz for a 700 MHz spectrometer which is consistent with the coupling of 8 Hz measured for EtG doublet in (Nicholas et al. 2006). To confirm this measurement on the aligned dataset, we measured this coupling on a raw NMR data of 15 single individuals for whom the pseudo-quantification of EtG is above 3 standard deviations from the mean EtG pseudo-quantification in the whole population (Supplementary Figure 23). The average doublet coupling constant measured ($\pm$standard deviation) among the 15 NMR samples is equal to $8.06 \pm 0.17$ Hz confirming the identification of EtG doublet in these individuals.

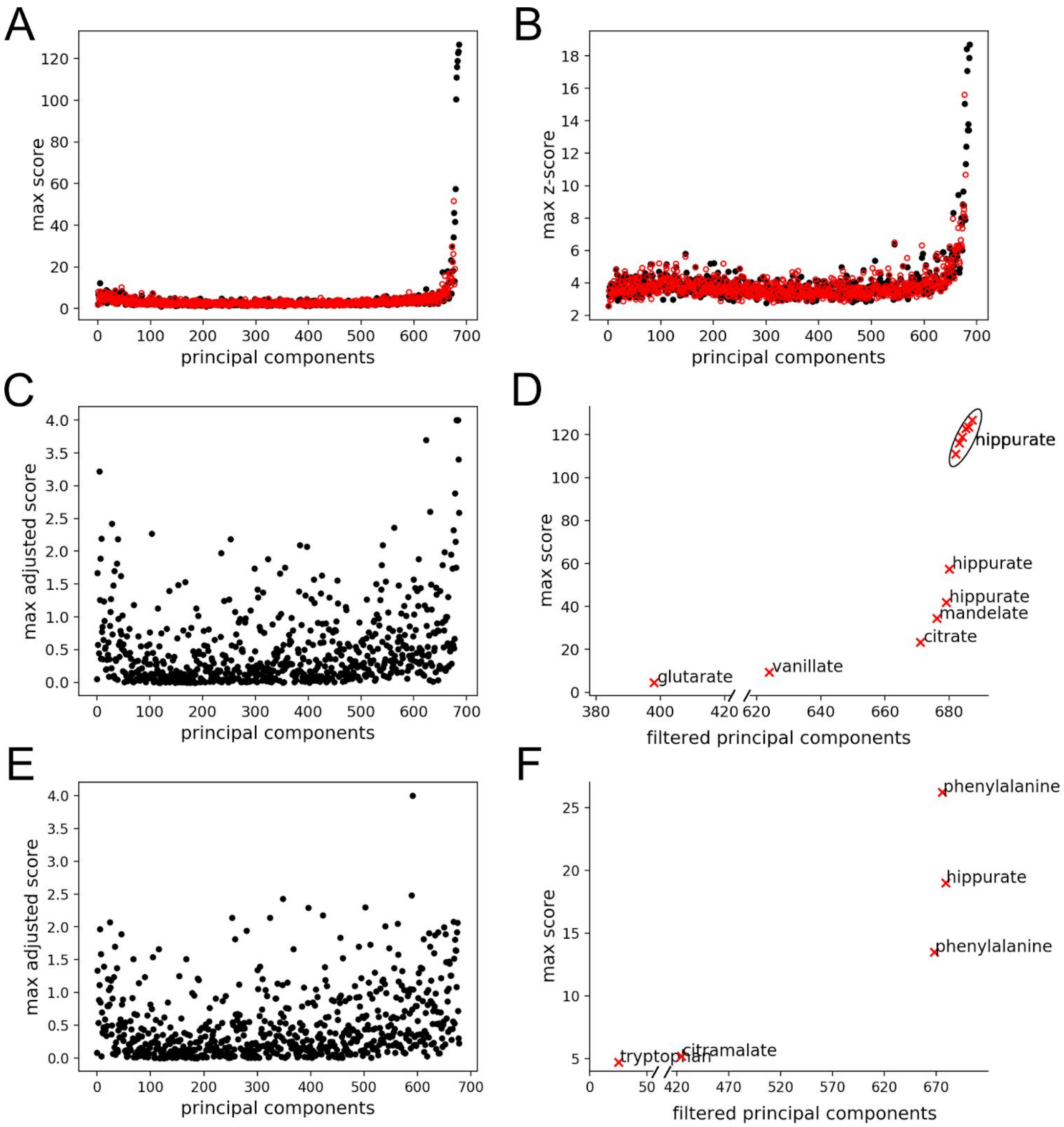


Supplementary Figure 1: Principal component pseudospectra analysis. (A) maximum metabomatching score (chi-squared mode) achieved for each principal components of the full metabolome (in black) and for principal components after removing the highly correlated features from the metabolome (in red). (B) maximum absolute value of z_score obtained through z-score transformation of each principal component of the full metabolome (in black) and for principal components after removing the highly correlated features from the metabolome (in red). (C) maximum metabomatching adjusted score achieved for each principal components for full metabolome. (D) maximum metabomatching score only for principal components that pass the filtering (adjusted score > 2 and max z-score > 4) for full metabolome and their matching metabolites. (E) maximum metabomatching adjusted score achieved for each principal components after removing the highly correlated features from the metabolome. (F) maximum metabomatching score only for principal components that pass the filtering (adjusted score > 2 and max z-score > 4) after removing the highly correlated features from the metabolome and their matching metabolites.

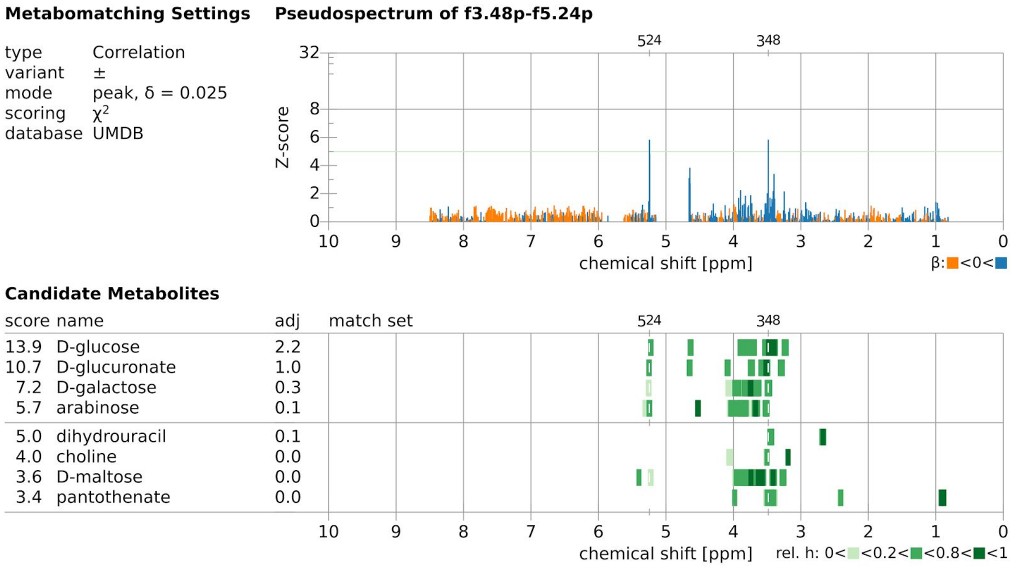


Supplementary Figure 2: Metabomatching for ACP module 3.48 & 5.24 matching D-glucose.

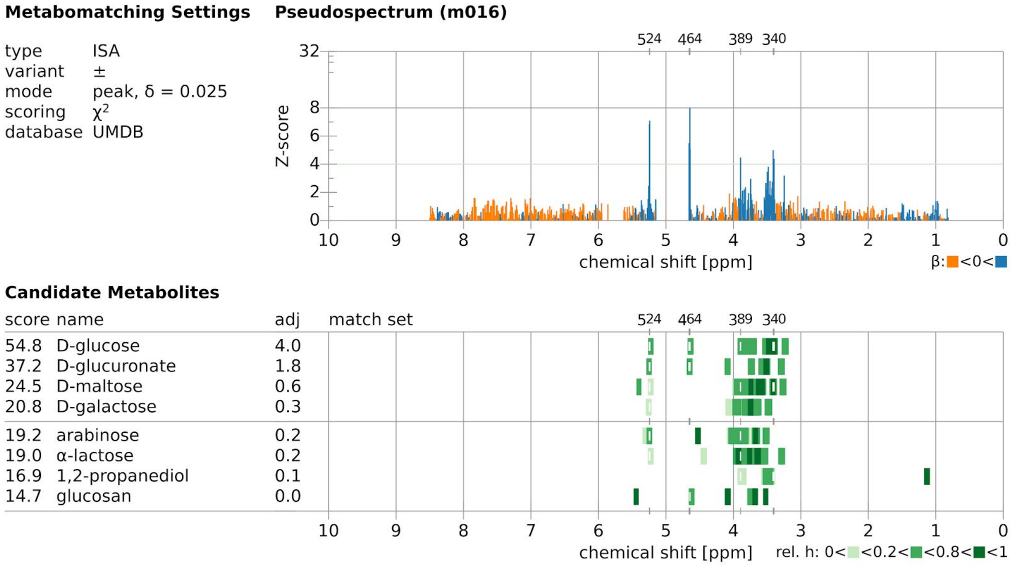


Supplementary Figure 3: Metabomatching for ISA module #016 matching D-glucose.

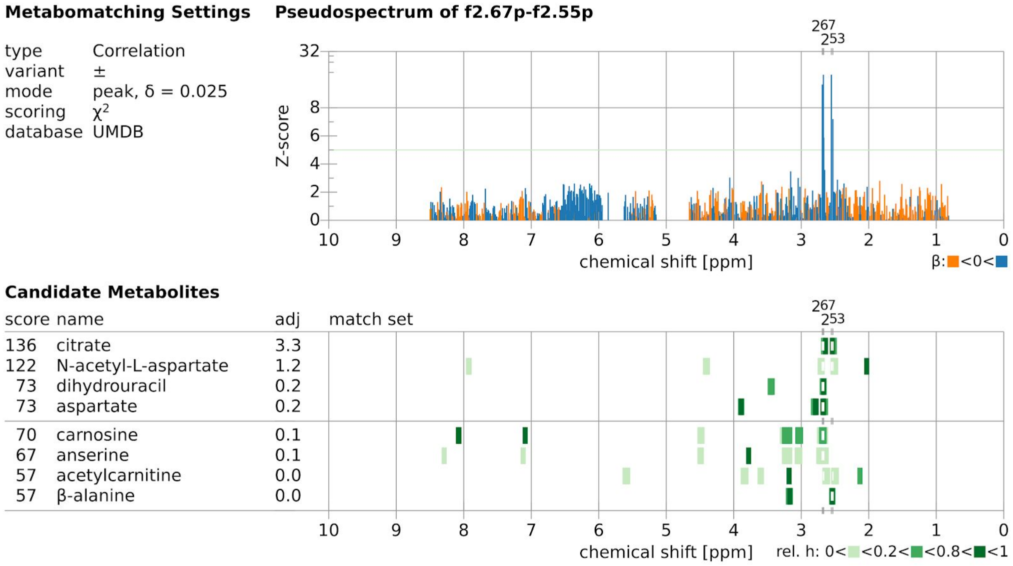


Supplementary Figure 4: Metabomatching for ACP module 2.67 & 2.55 matching citrate.

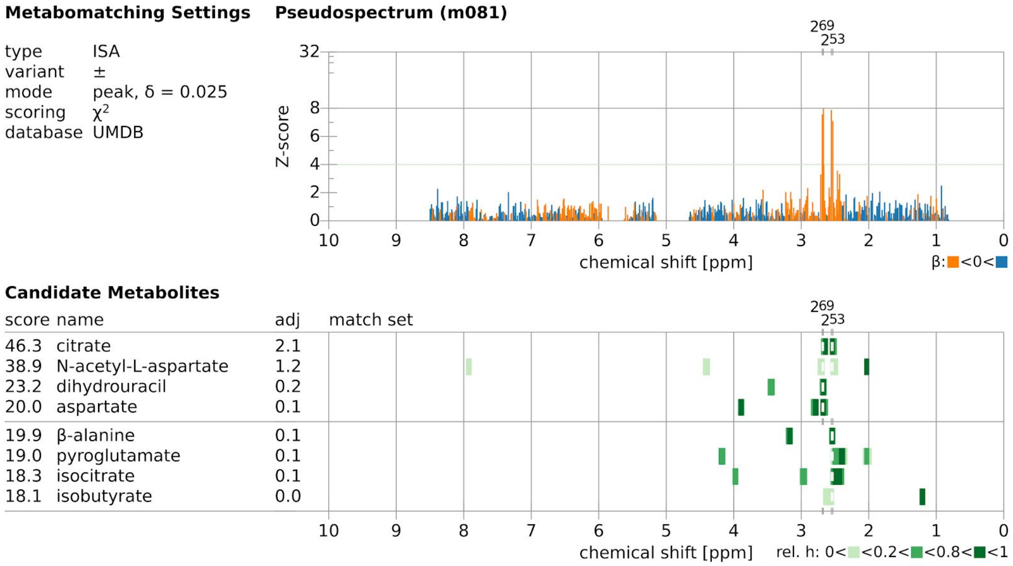


Supplementary Figure 5: Metabomatching for ISA module #081 matching citrate.

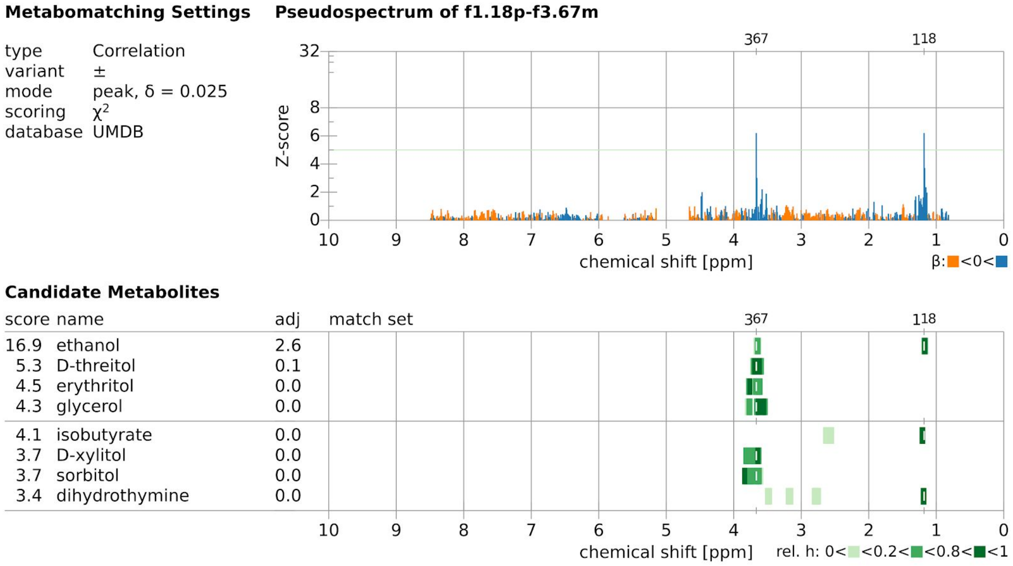


Supplementary Figure 6: Metabomatching for ACP module 1.18 & 3.67 matching ethanol.

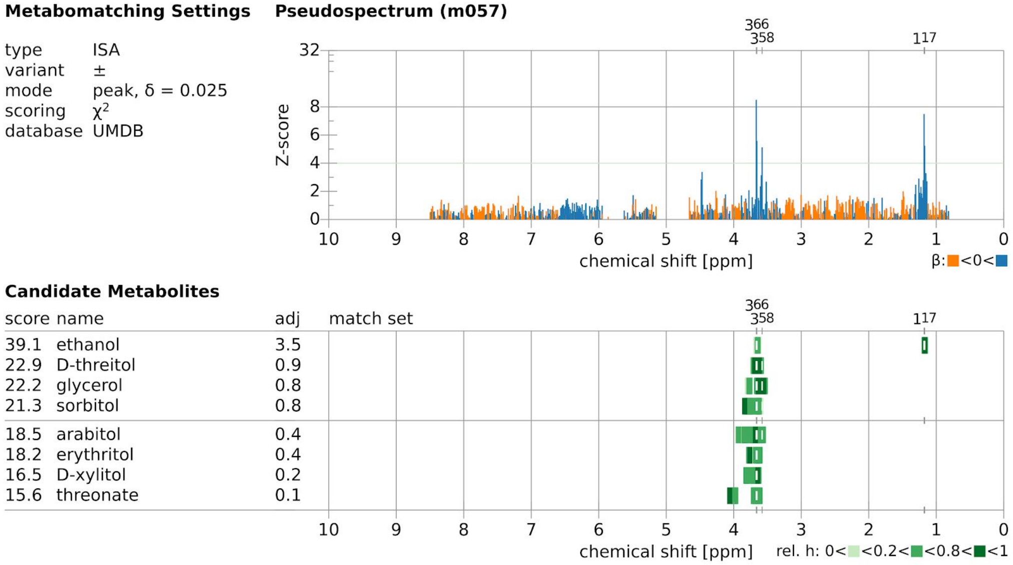


Supplementary Figure 7: Metabomatching for ISA module #057 matching ethanol.

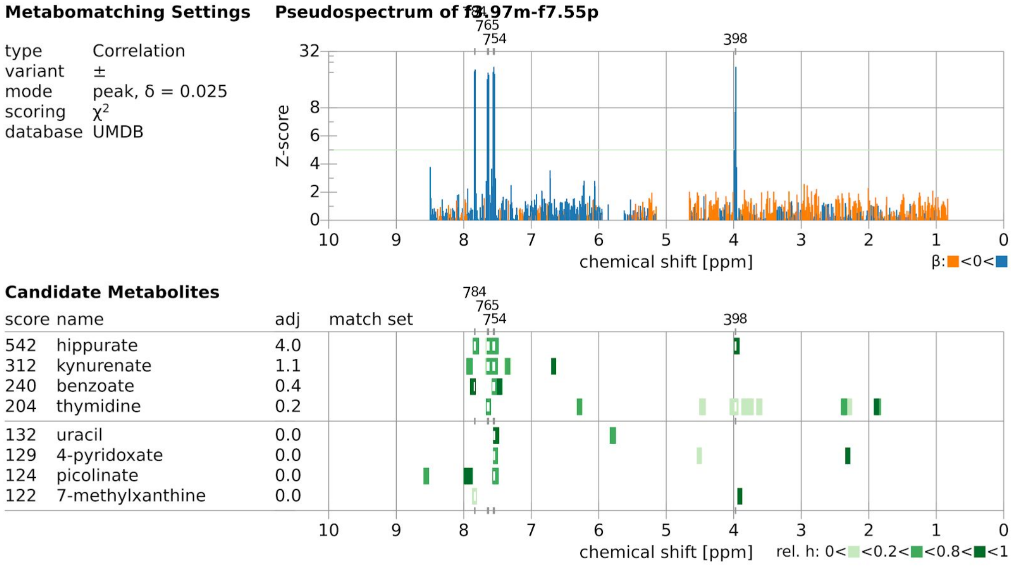


Supplementary Figure 8: Metabomatching for ACP module 3.97 & 7.55 matching hippurate.

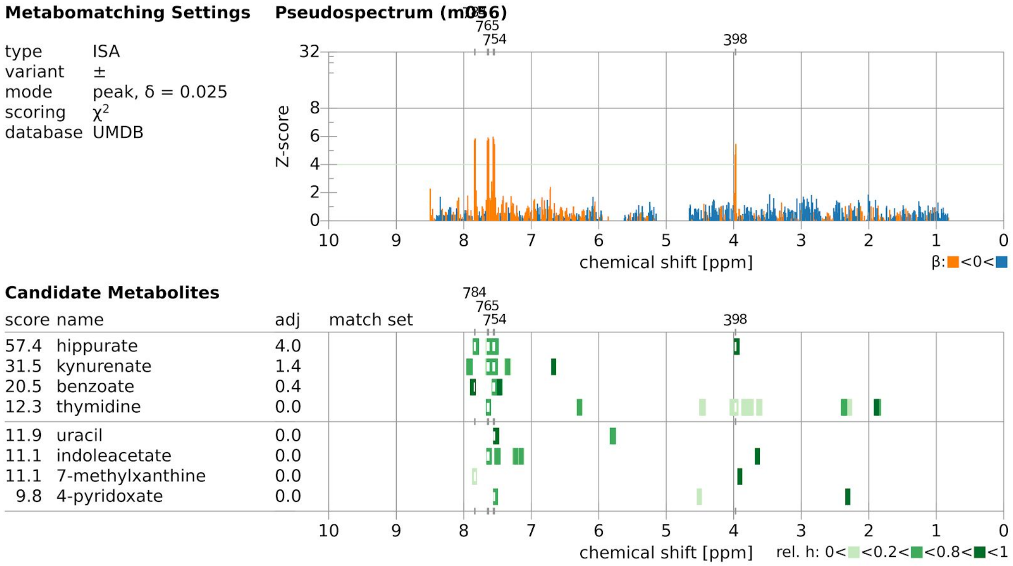


Supplementary Figure 9: Metabomatching for ISA module #056 matching hippurate.

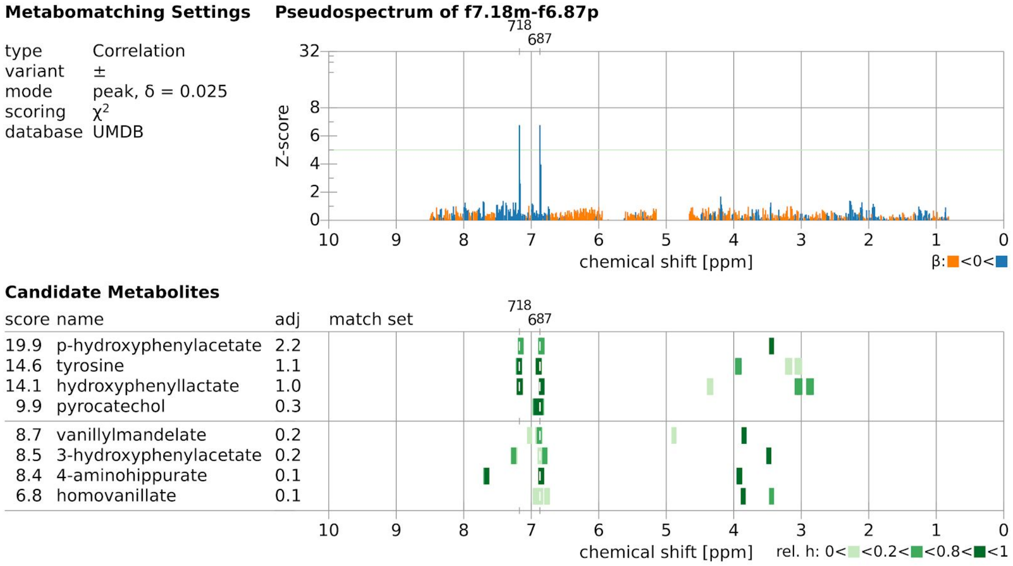


Supplementary Figure 10: Metabomatching for ACP module 7.18 & 6.87 matching p-hydroxyphenylacetate.

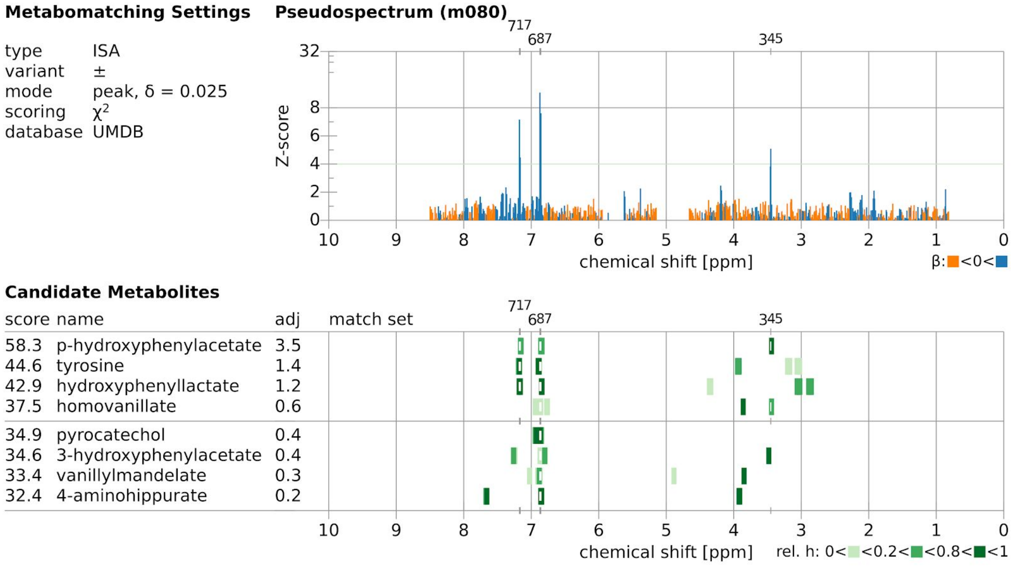


Supplementary Figure 11: Metabomatching for ISA module #080 matching p-hydroxyphenylacetate.

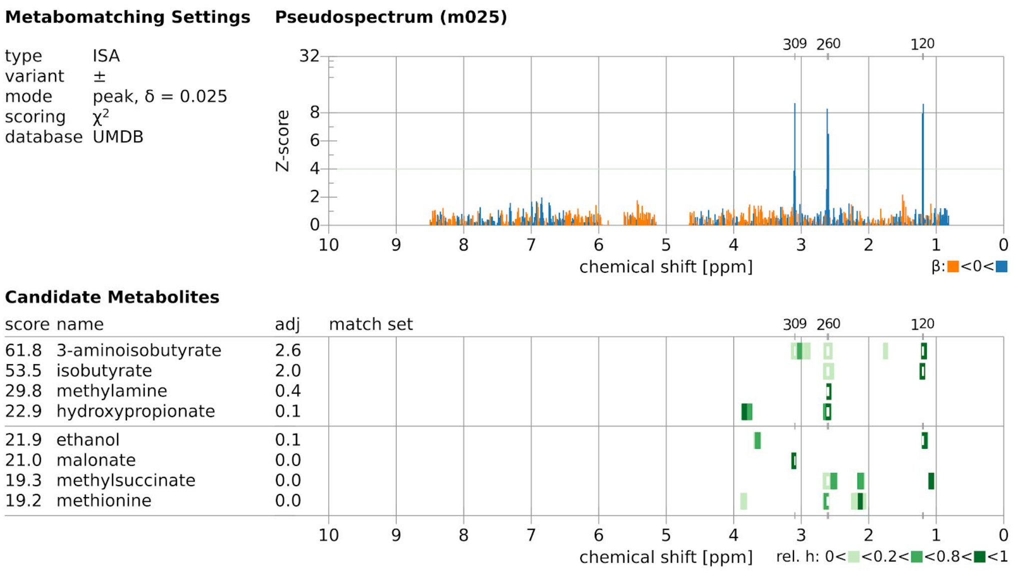


Supplementary Figure 12: Metabomatching for ISA module #025 matching 3-aminoisobutyrate.

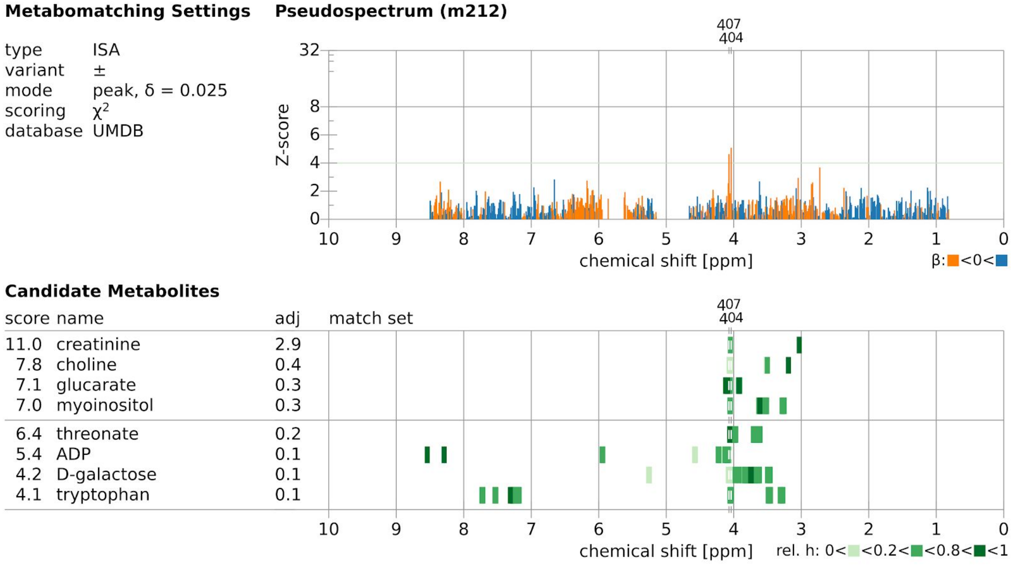


Supplementary Figure 13: Metabomatching for ISA module #212 matching creatinine.

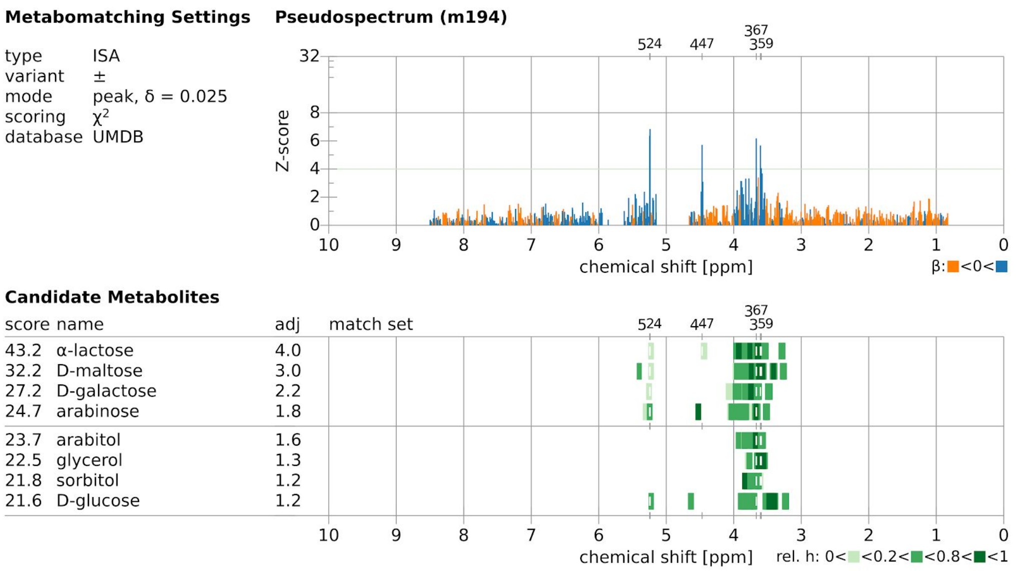


Supplementary Figure 14: Metabomatching for ISA module #194 matching alpha-lactose.

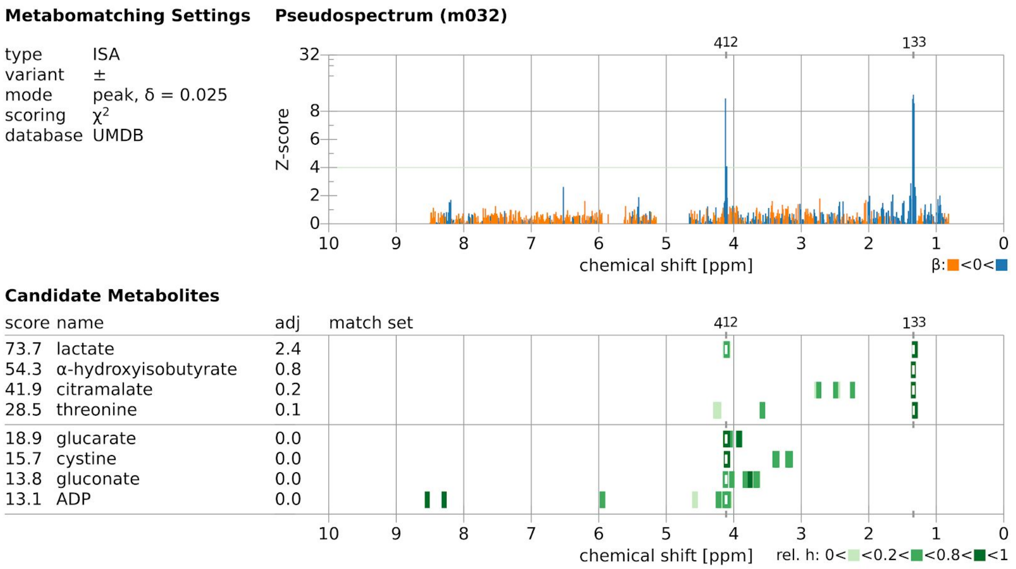


Supplementary Figure 15: Metabomatching for ISA module #032 matching lactate.

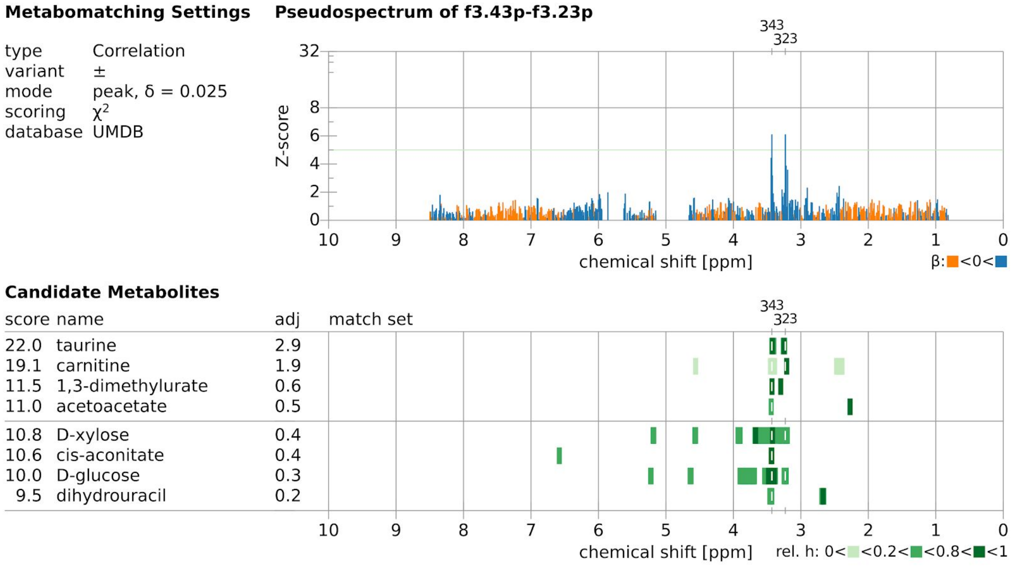


Supplementary Figure 16: Metabomatching for ACP module 3.43 & 3.23 matching taurine.

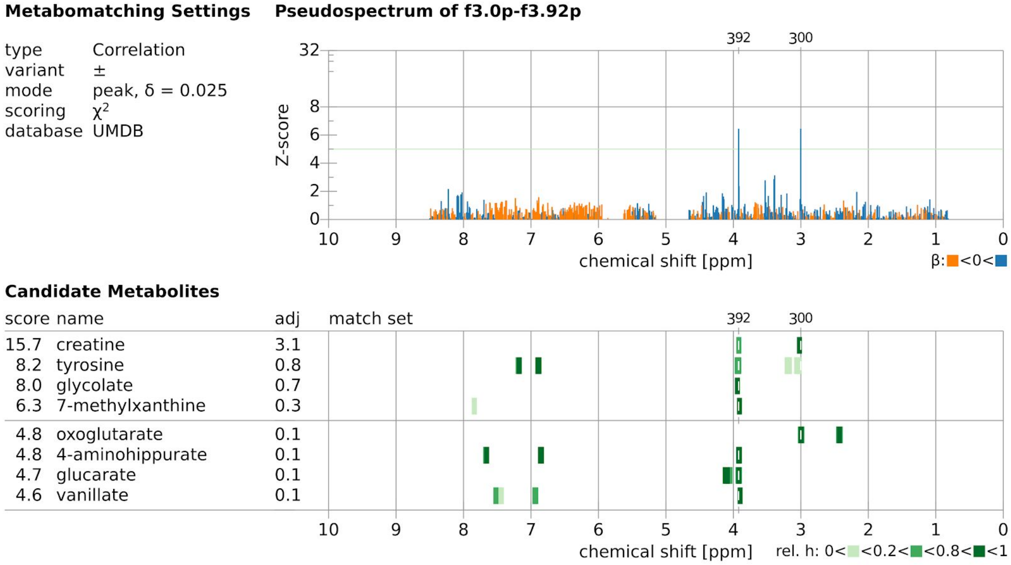


Supplementary Figure 17: Metabomatching for ACP module 3.00 & 3.92 matching creatine.

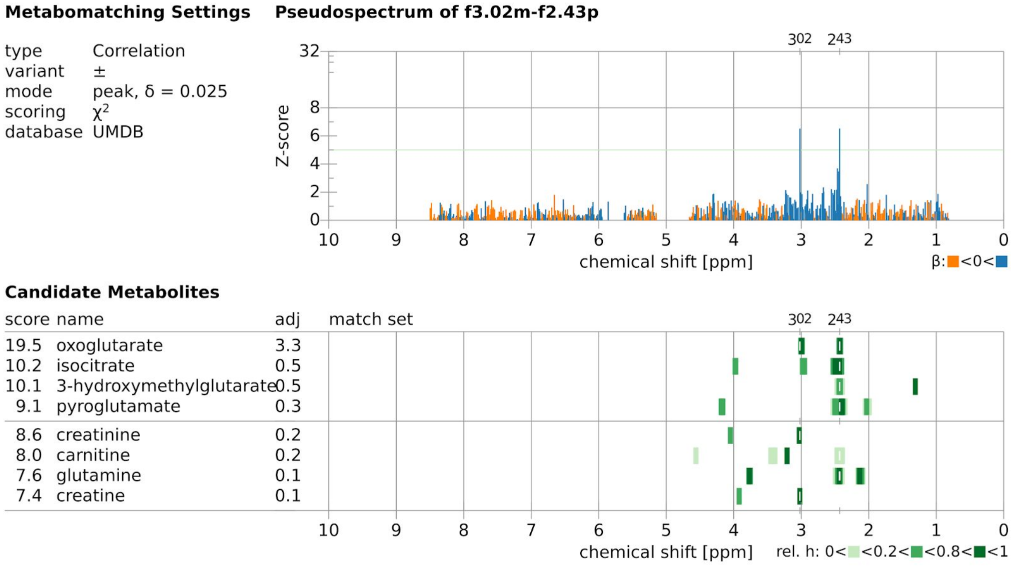


Supplementary Figure 18: Metabomatching for ACP module 3.02 & 2.43 matching oxoglutarate.

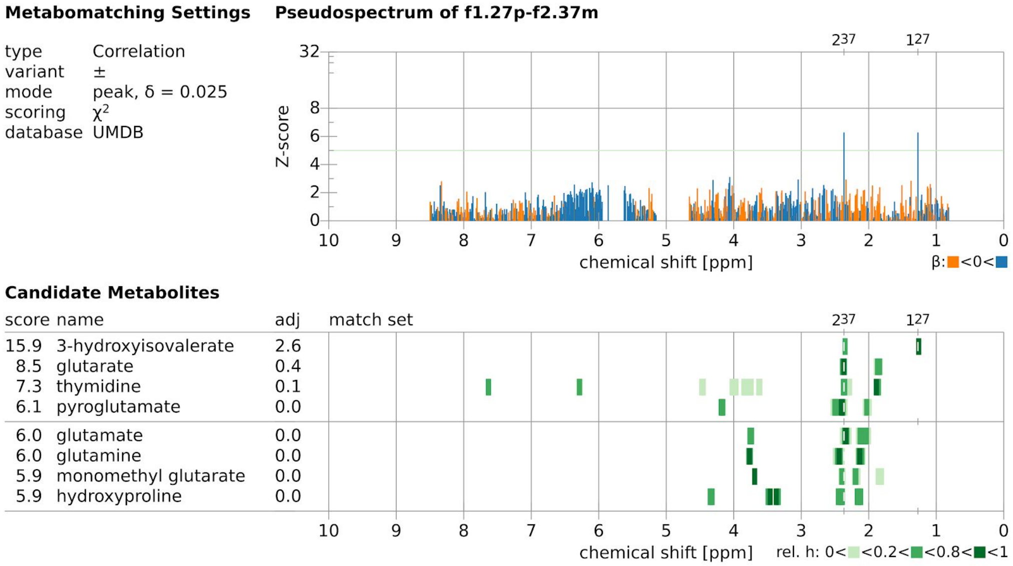


Supplementary Figure 19: Metabomatching for ACP module 1.27 & 2.37 matching 3-hydroxyisovalerate.

| features (ppm) | correlation with CDT |
|---|---|
| 1.202 | 0.11 |
| 1.191 | 0.14 |
| 1.180 | 0.26 |
| 1.172 | 0.11 |
| 1.164 | -0.04 |
| 1.163 | 0.11 |
| 1.153 | 0.45 |
| 1.149 | 0.35 |
| 1.140 | 0.44 |
| 1.130 | 0.07 |

Supplementary Figure 20: Correlation between features 1.13-1.20 and serum CDT .

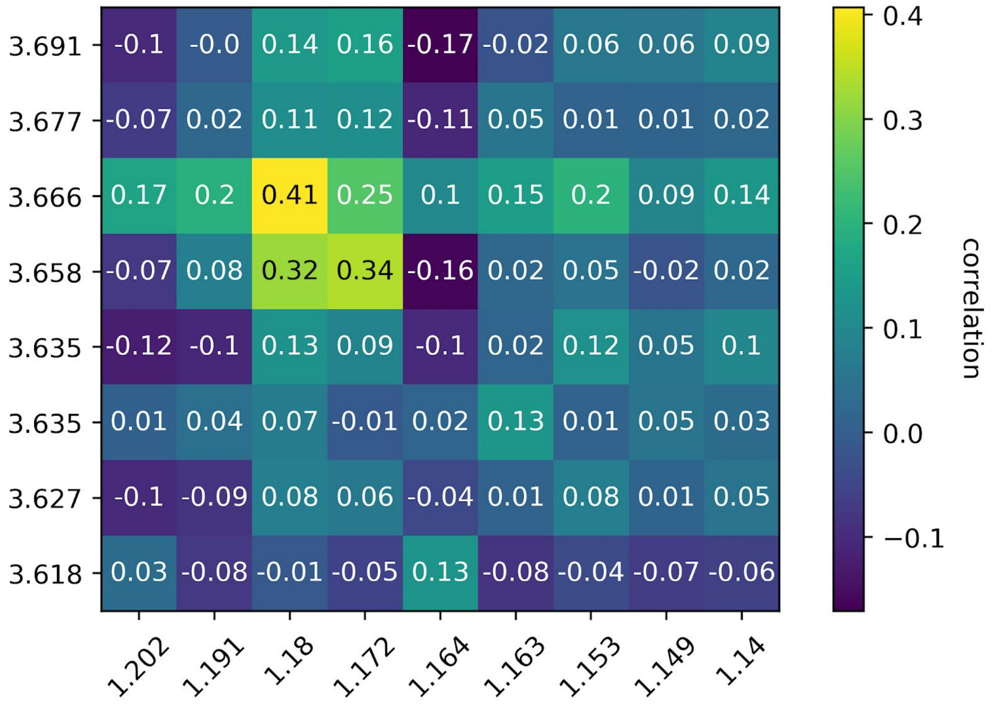


Supplementary Figure 21: Correlation between features 1.14-1.20 ppm and 3.62-3.69 ppm, the regions of ethanol's two multiplets based on UMDB.

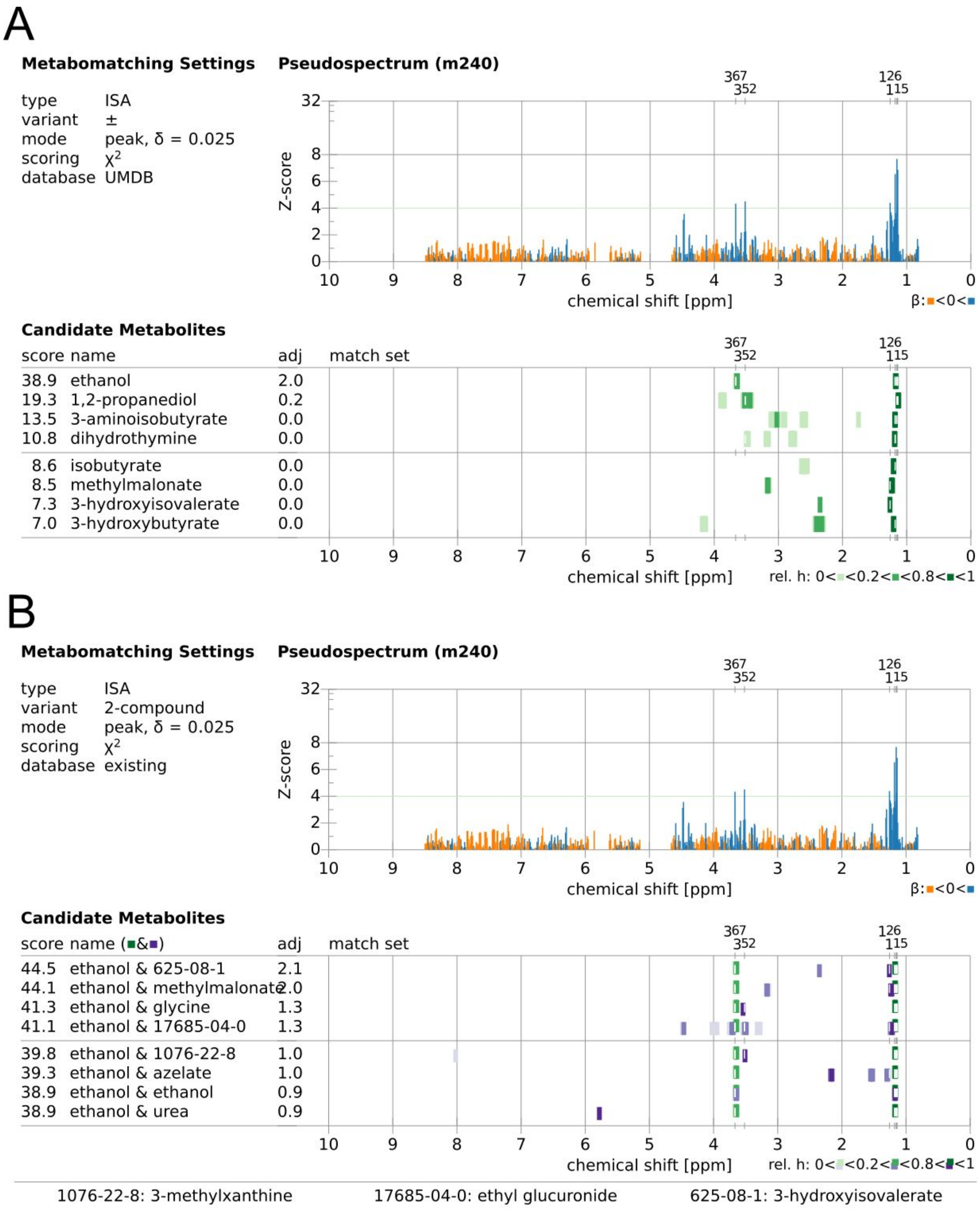


Supplementary Figure 22: metabomatching figures of ISA module #240 (A) one compound search and (B) two compound search. Ethanol and EtG together are a better match for the pseudospectra rather than ethanol alone.

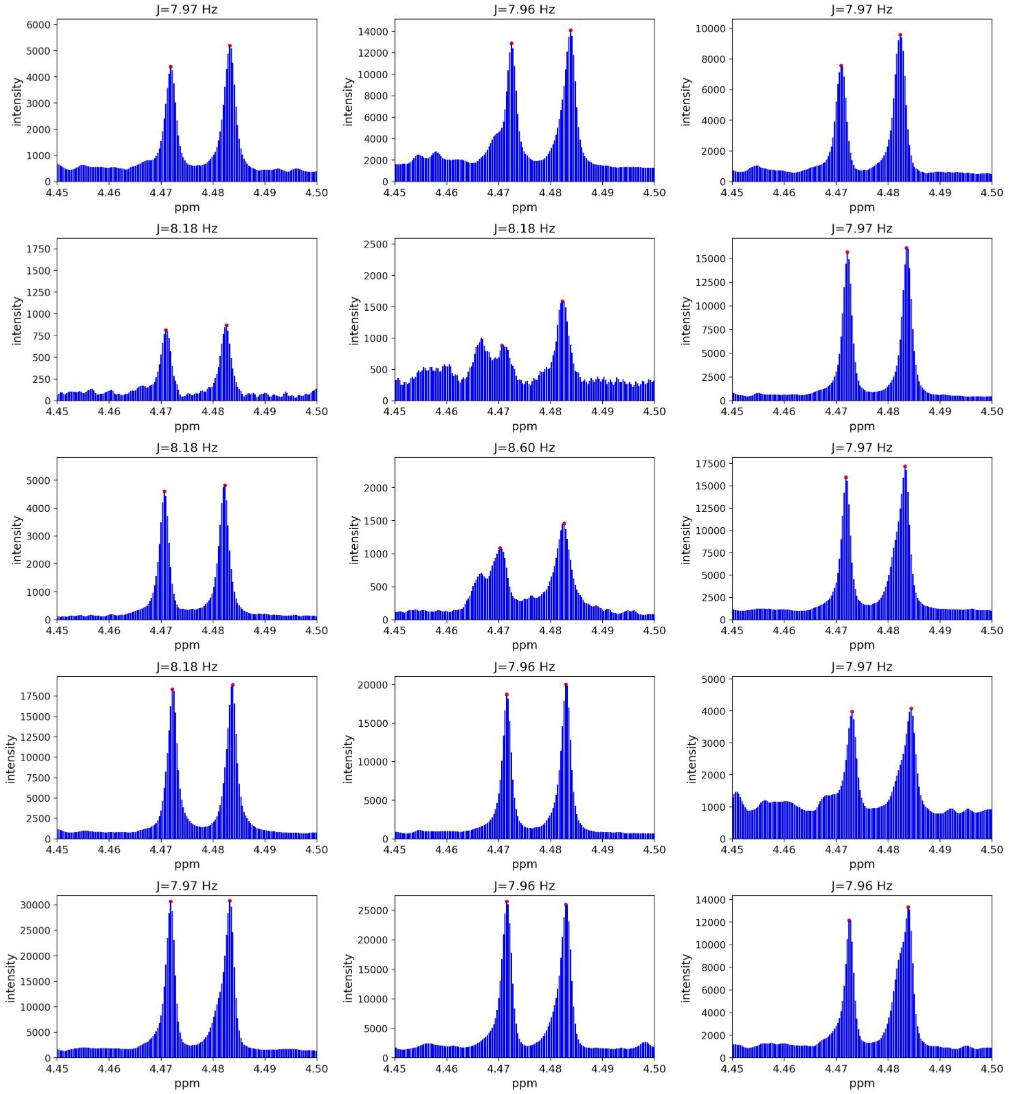


Supplementary Figure 23: The raw NMR spectra of 15 individuals with EtG pseudo-quantification above 3 standard deviation from the population mean shown in the region of 4.45 - 4.50 ppm, to measure EtG doublet coupling constant. The two peaks of the doublet are highlighted by red dots. The average calculated coupling constant ($\pm$ standard deviation) is $8.06 \pm 0.17$ Hz.

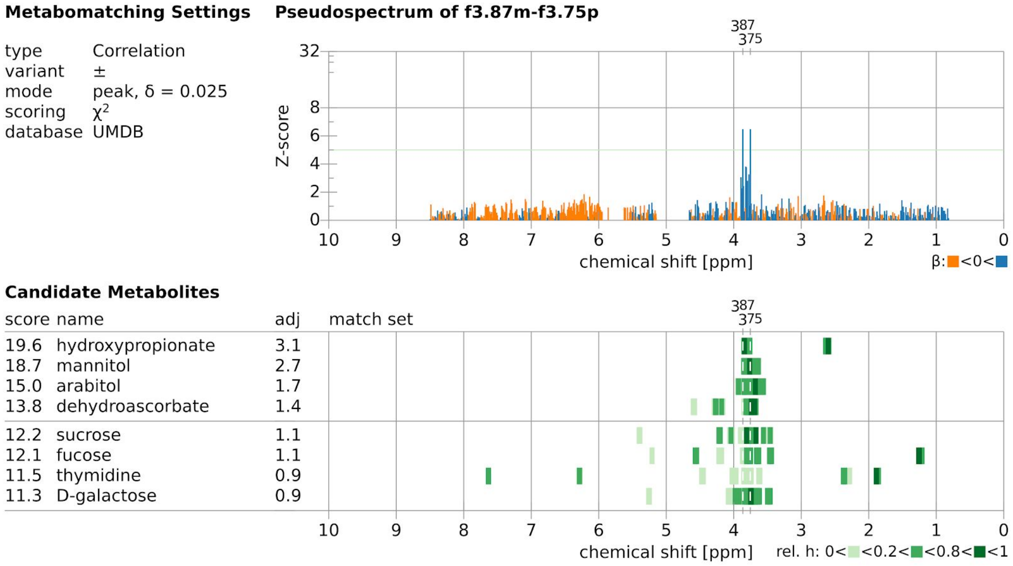


Supplementary Figure 24: Metabomatching for ACP module 3.87 & 3.75 matching hydroxypropionate.

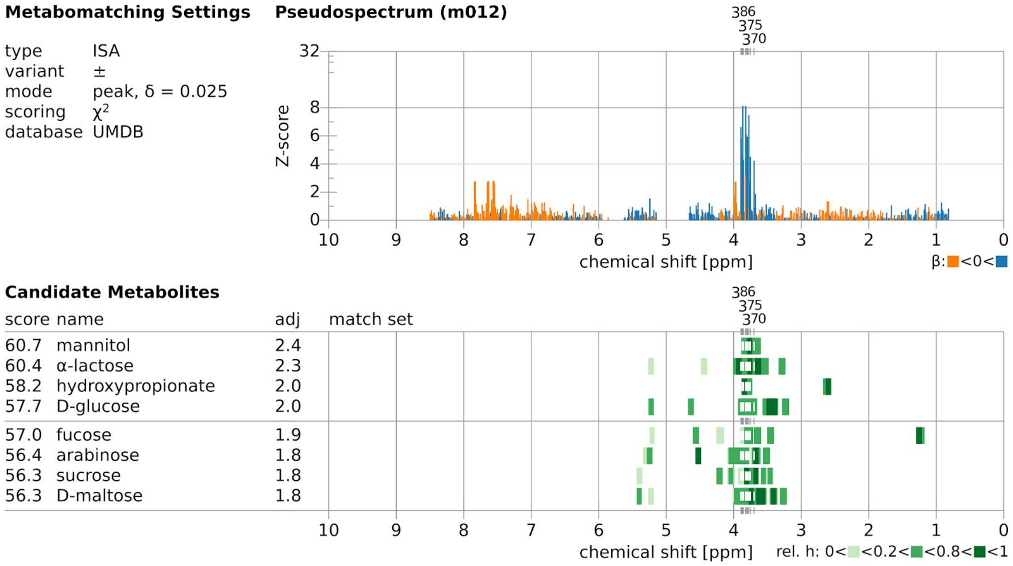


Supplementary Figure 25: Metabomatching for ISA module #012 matching mannitol.

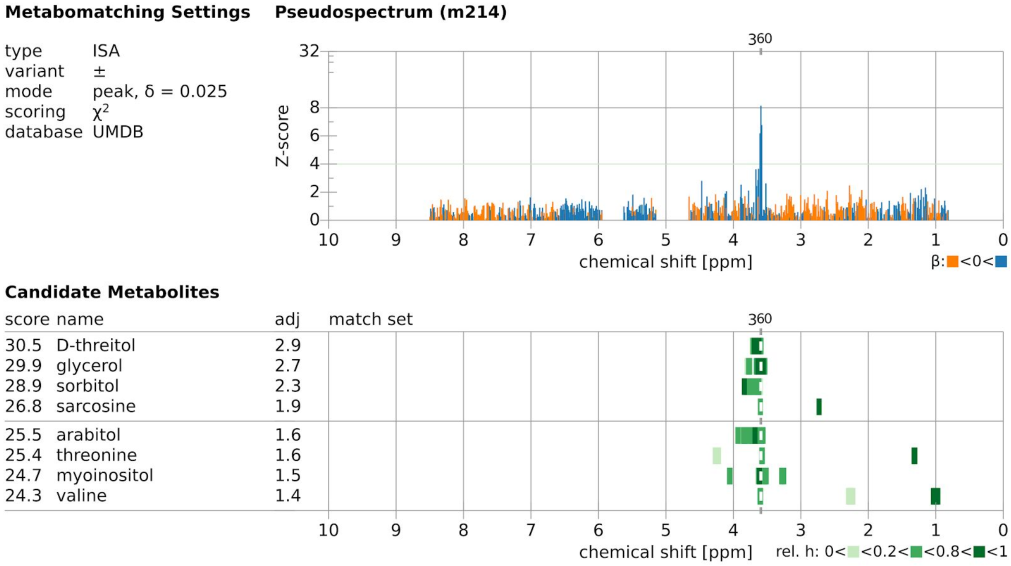


Supplementary Figure 26: Metabomatching for ISA module #214 matching D-threitol.

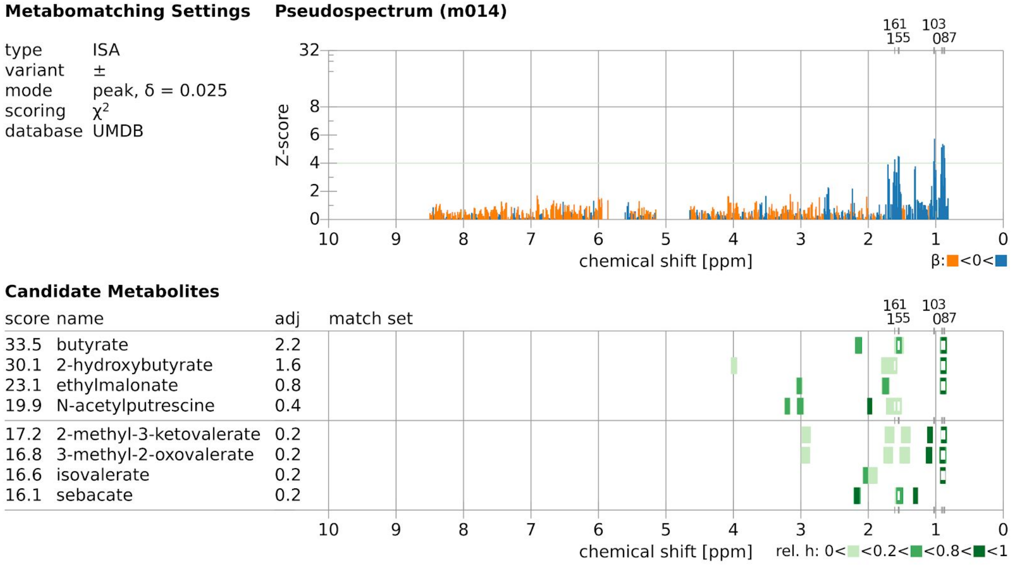


Supplementary Figure 27: Metabomatching for ISA module #014 matching butyrate.

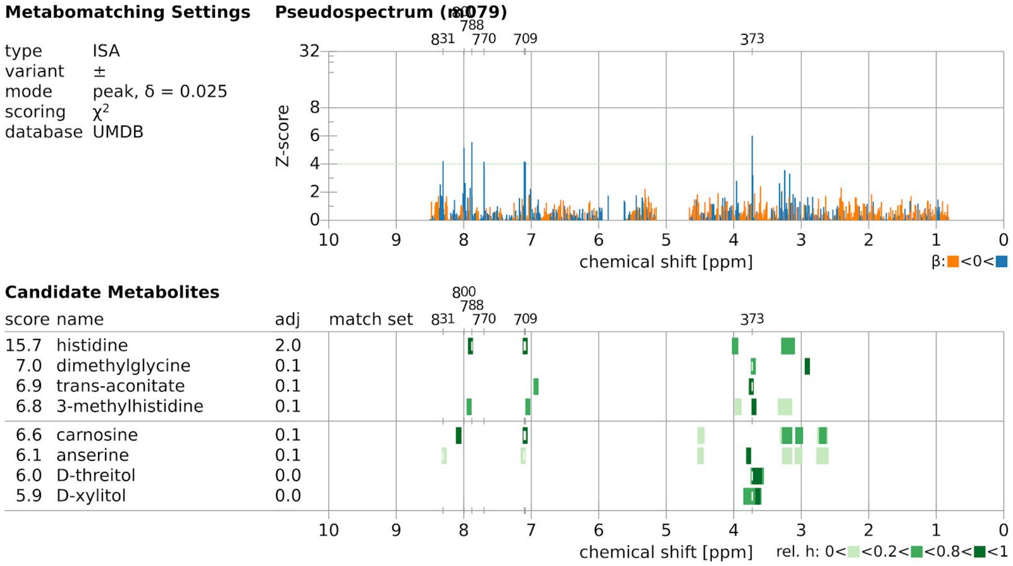


Supplementary Figure 28: Metabomatching for ISA module #079 matching histidine.

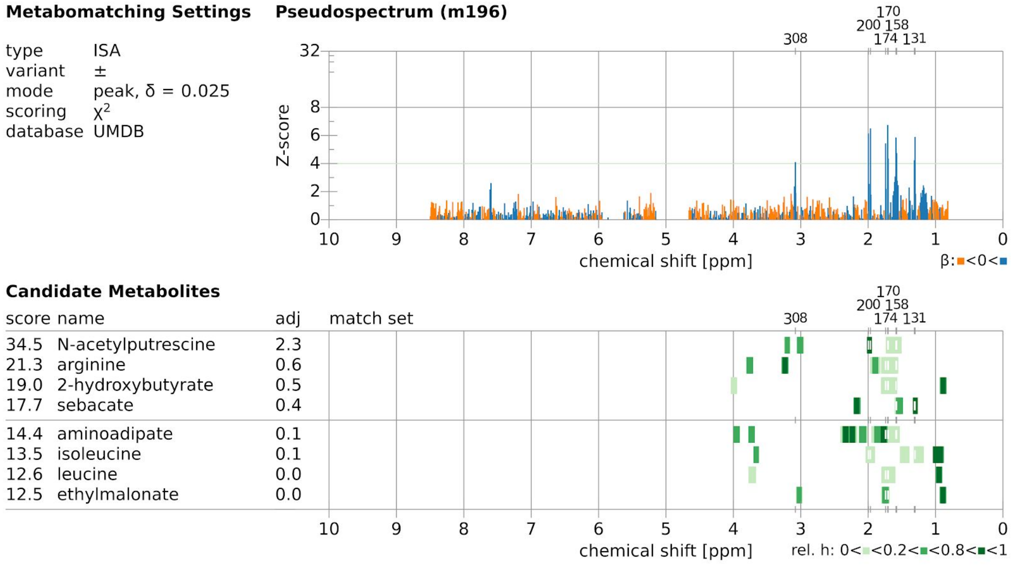


Supplementary Figure 29: Metabomatching for ISA module #196 matching N-acetylputrescine.

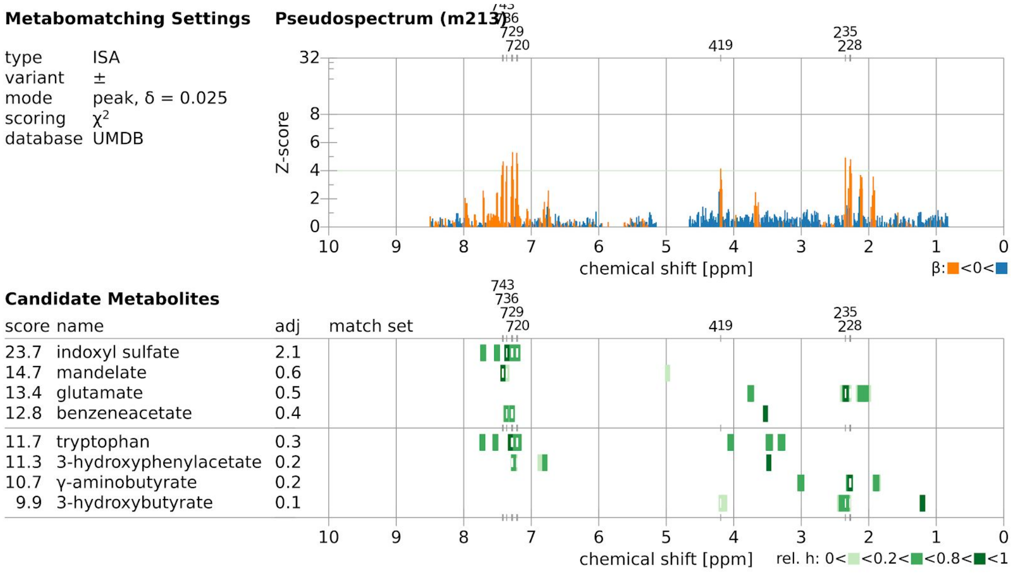


Supplementary Figure 30: Metabomatching for ISA module #213 matching indoxyl sulfate.

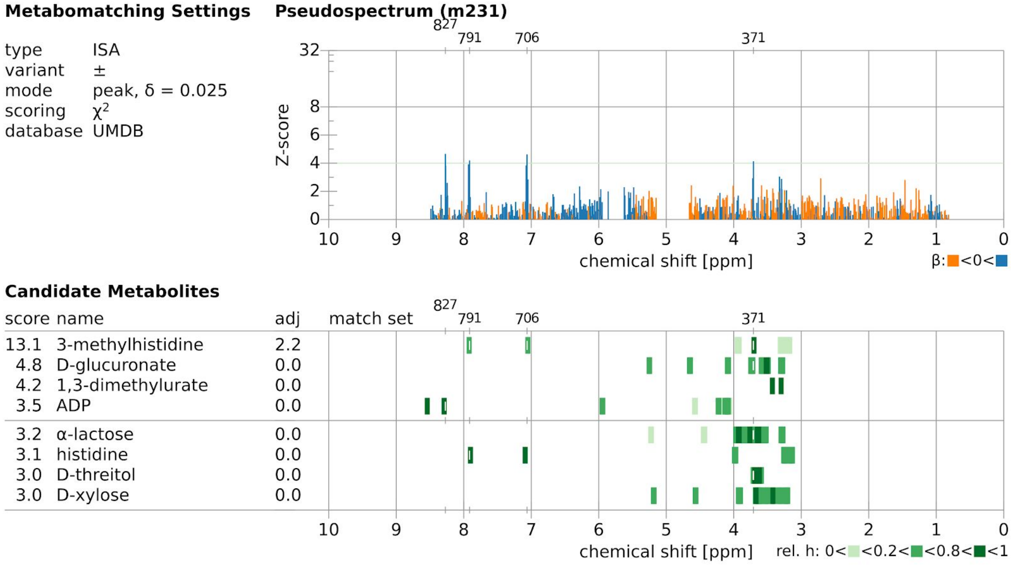


Supplementary Figure 31: Metabomatching for ISA module #231 matching 3-methylhistidine.

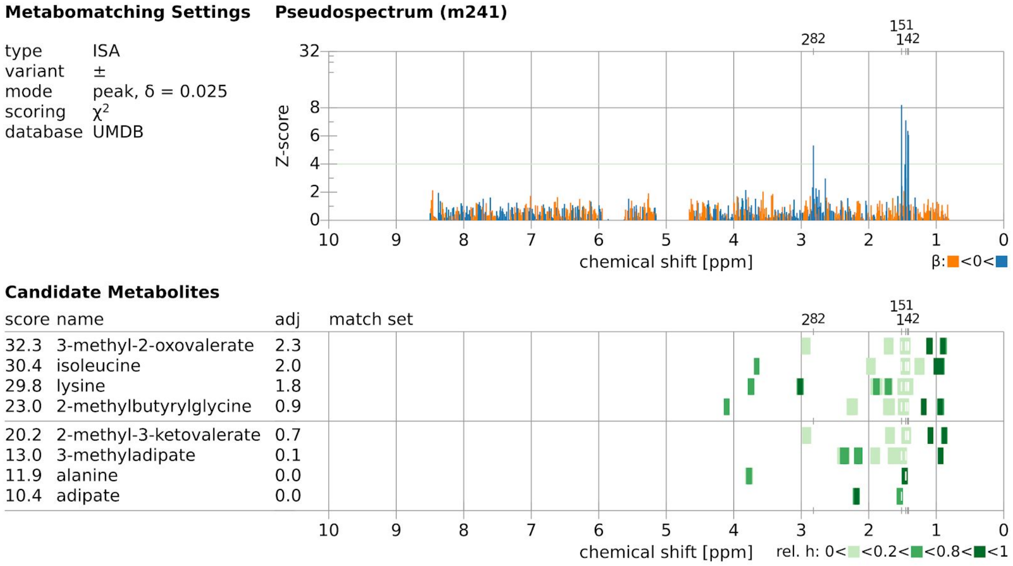


Supplementary Figure 32: Metabomatching for ISA module #241 matching 3-methyl-2-oxovalerate.

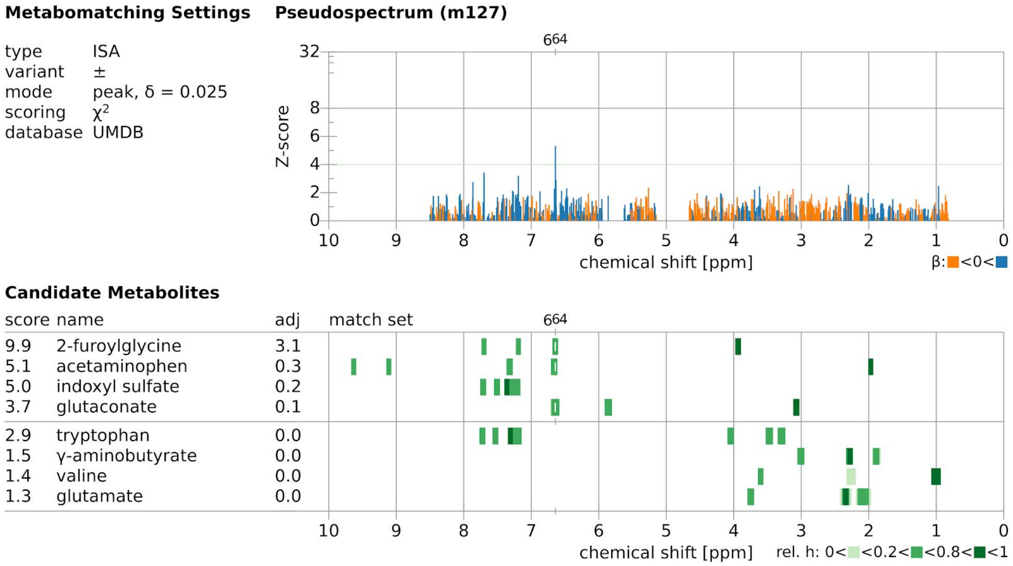


Supplementary Figure 33: Metabomatching for ISA module #127 matching 2-furoylglycine.

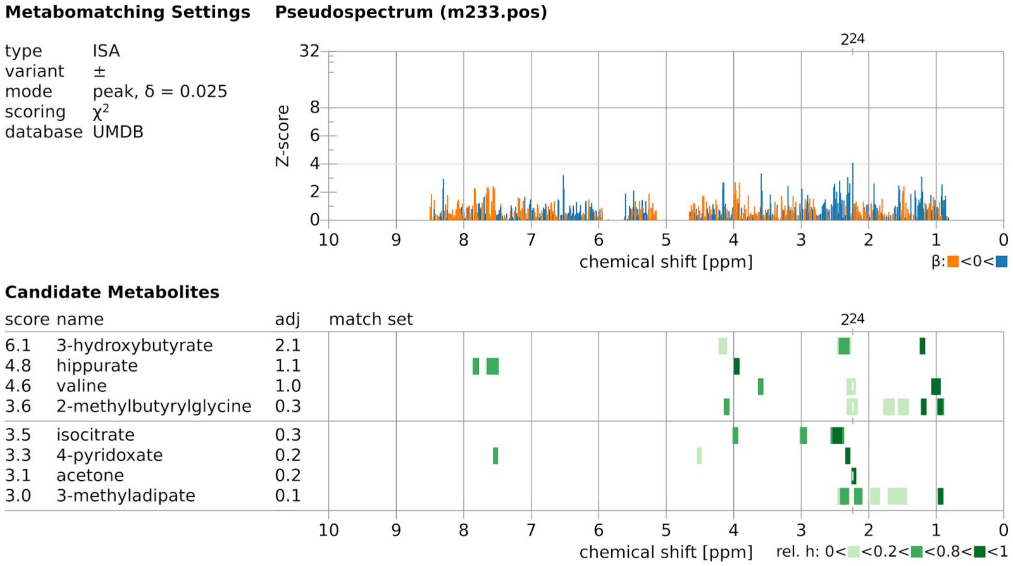


Supplementary Figure 34: Metabomatching for ISA module #233 matching 3-hydroxybutyrate.

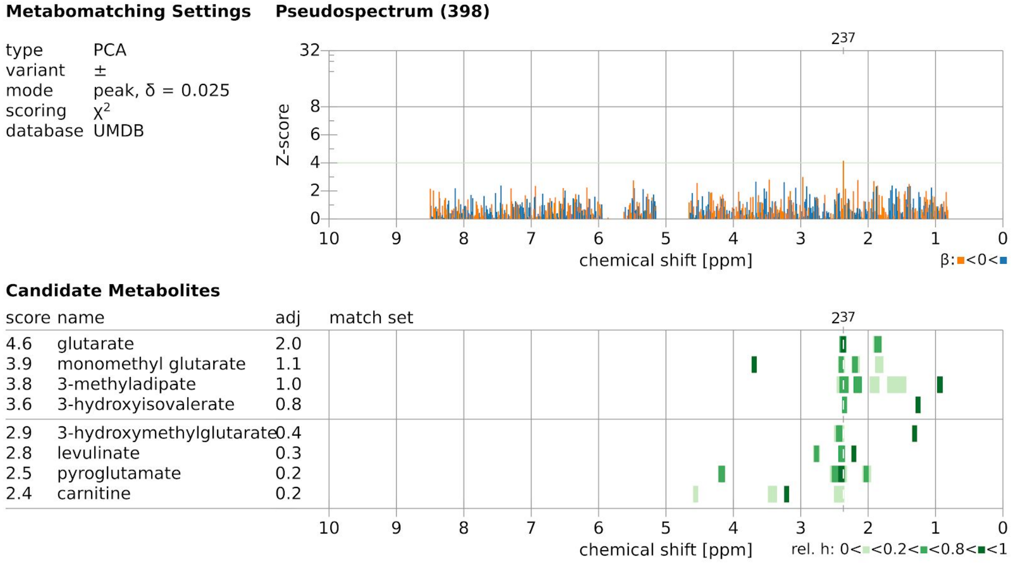


Supplementary Figure 35: Metabomatching for principal component module #398 matching glutarate.

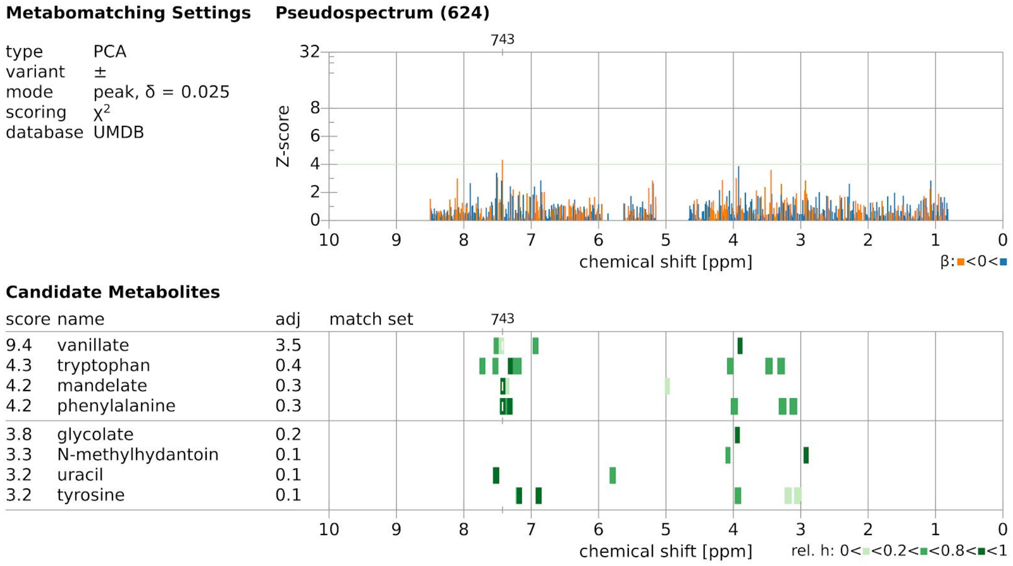


Supplementary Figure 36: Metabomatching for principal component module #624 matching vanillate.

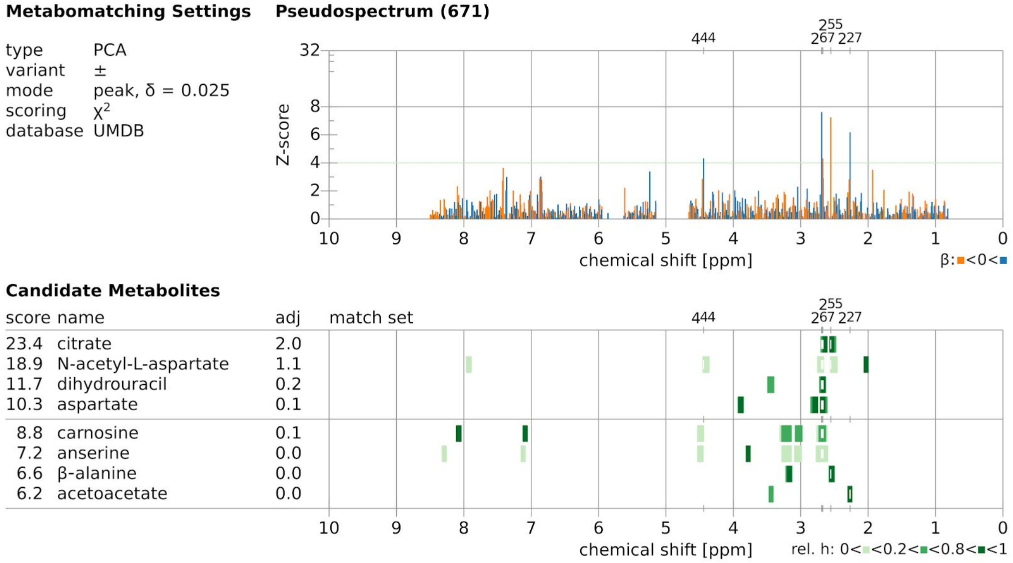


Supplementary Figure 37: Metabomatching for principal component #671 matching citrate.

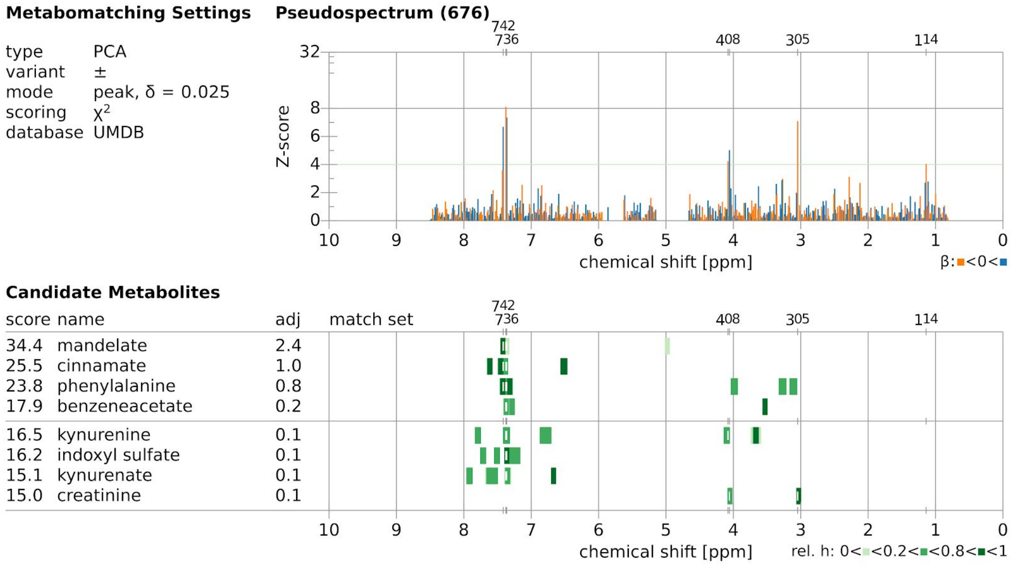


Supplementary Figure 38: Metabomatching for principal component module #676 matching mandelate.

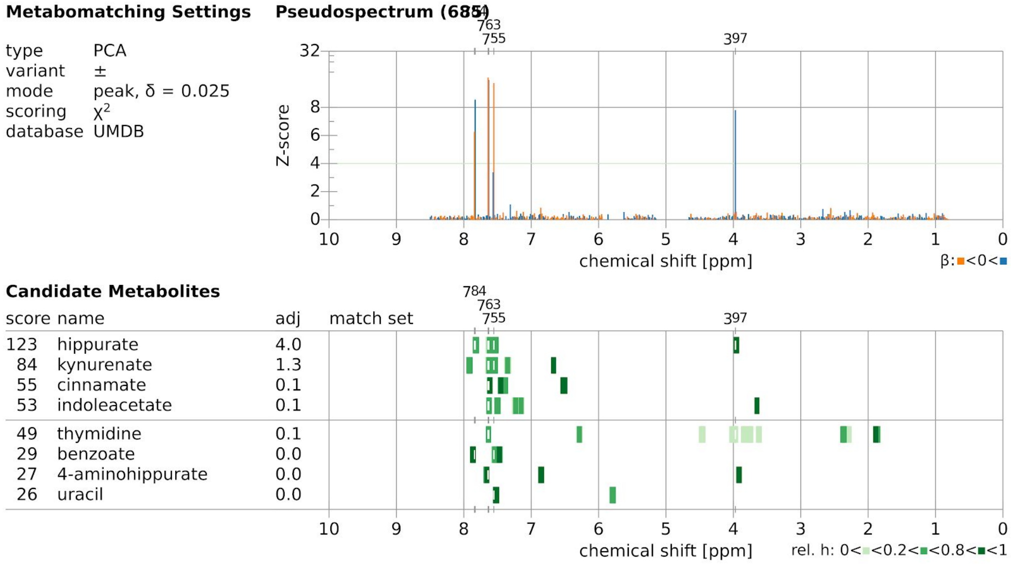


Supplementary Figure 39: Metabomatching for principal component #685 matching hippurate.

Supplementary Table 1: Correlation between pseudo-quantification and measured biomarkers of ethanol, serum gamma-glutamyl transferase (GGT), asialotransferrin (ATRN) and self-reported alcohol consumption.

| **Urine Metabolite** | **Feature source** | **Multiplet positions** | **Related bio-marker** | **Correlation with 95% CI** |
|---|---|---|---|---|
| ethanol | UMDB | 1.17, 3.65 | Serum GGT | 0.10 [0.04, 0.16] |
| ethanol | UMDB | 1.17, 3.65 | Serum ATRN | 0.21 [0.15, 0.27] |
| ethanol | UMDB | 1.17, 3.65 | Self report | 0.29 [0.23, 0.35] |
| ethanol | ACP: f1.18-f3.67 ISA: Module #57 | 1.18, 3.67 | Serum GGT | 0.00 [-0.06, 0.06] |
| ethanol | ACP: f1.18-f3.67 ISA: Module #57 | 1.18, 3.67 | Serum ATRN | 0.15 [0.08, 0.21] |
| ethanol | ACP: f1.18-f3.67 ISA: Module #57 | 1.18, 3.67 | Self report | 0.10 [0.03, 0.16] |
| EtG | (Nicholas et al. 2006) | 1.24, 3.30, 3.52, 3.71, 3.99, 4.48 | Serum GGT | 0.15 [0.09, 0.21] |
| EtG | (Nicholas et al. 2006) | 1.24, 3.30, 3.52, 3.71, 3.99, 4.48 | Serum ATRN | 0.17 [0.11, 0.23] |
| EtG | (Nicholas et al. 2006) | 1.24, 3.30, 3.52, 3.71, 3.99, 4.48 | Self report | 0.39 [0.33, 0.44] |
| EtG | ISA: Module #240 | 1.24, 3.52, 4.47 | Serum GGT | 0.21 [0.15, 0.27] |
| EtG | ISA: Module #240 | 1.24, 3.52, 4.47 | Serum ATRN | 0.25 [0.19, 0.31] |
| EtG | ISA: Module #240 | 1.24, 3.52, 4.47 | Self report | 0.48 [0.43, 0.54] |
